# Connectivity patterns that shape olfactory representation in a mushroom body network model

**DOI:** 10.1101/2021.02.10.430647

**Authors:** Daniel Zavitz, Elom A. Amematsro, Alla Borisyuk, Sophie J.C. Caron

## Abstract

Cerebellum-like structures — such as the insect mushroom body — are found in many brains and share a basic fan-out–fan-in network architecture. How the specific structural features of these networks give rise to their learning function remains largely unknown. To investigate this structure–function relationship, we developed a minimal computational model of the extensively studied *Drosophila melanogaster* mushroom body. We show how well-defined connectivity patterns between the Kenyon cells — the constituent neurons of the mushroom body — and their input projection neurons endow different functions, enabling the mushroom body to process olfactory information more efficiently. First, biases in the likelihoods at which individual projection neurons connect to Kenyon cells allow the mushroom body to prioritize the learning of particular, ethologically meaningful odors. Second, groups of projection neurons connecting preferentially to the same Kenyon cells facilitate the mushroom body to generalize across similar odors. Altogether, our results demonstrate how different connectivity patterns shape the representation space of a well-studied cerebellum-like network and impact its learning outcomes.

## INTRODUCTION

Cerebellum-like structures — such as the vertebrate cerebellum and the insect mushroom body are found in different brain regions of many different animal groups. These structures share a similar network architecture that supports different forms of associative learning (Bell et al., 2008; Farris, 2011). In short, cerebellum-like structures are formed by a large number of encoding neurons that receive input from a small number of input neurons and connect to a small number of output neurons, forming a typical fan-out–fan-in network (Figure 1A). In all cerebellum-like structures, sensory information is represented by the encoding neurons, whereas the output neurons drive different learned behavioral outcomes. Upon learning, the strength of the connections between encoding neurons and output neurons is modified such that specific associations can be formed (Owald and Waddell, 2015; Sawtell and Bell, 2008). Because of their well-characterized connectivity and functional features, cerebellum-like structures have been the focus of many theoretical studies that have fundamentally shaped our understanding of how associative learning is implemented at the level of a neuronal network. For instance, theoretical models predicted that the divergence of a small number of input neurons onto a large number of encoding neurons enables cerebellum-like structures to represent stimuli as sparse and largely non-overlapping activity patterns (Albus, 1971; Jortner et al., 2007; Marr, 1969). Such activity patterns are thought to be advantageous for learning, because they enable encoding neurons to represent a large number of stimuli as unique activity patterns, thus maximizing the representation space generated by a network (Cayco-Gajic et al., 2017; Litwin-Kumar et al., 2017; Stevens, 2015). Several theoretical studies have demonstrated that purely random connections between input neurons and encoding neurons allow networks to attain the largest possible representation space (Albus, 1971; Babadi and Sompolinsky, 2014; Cayco-Gajic and Silver, 2019; Jortner et al., 2007; Litwin-Kumar et al., 2017; Stevens, 2015).

**Figure 1.**
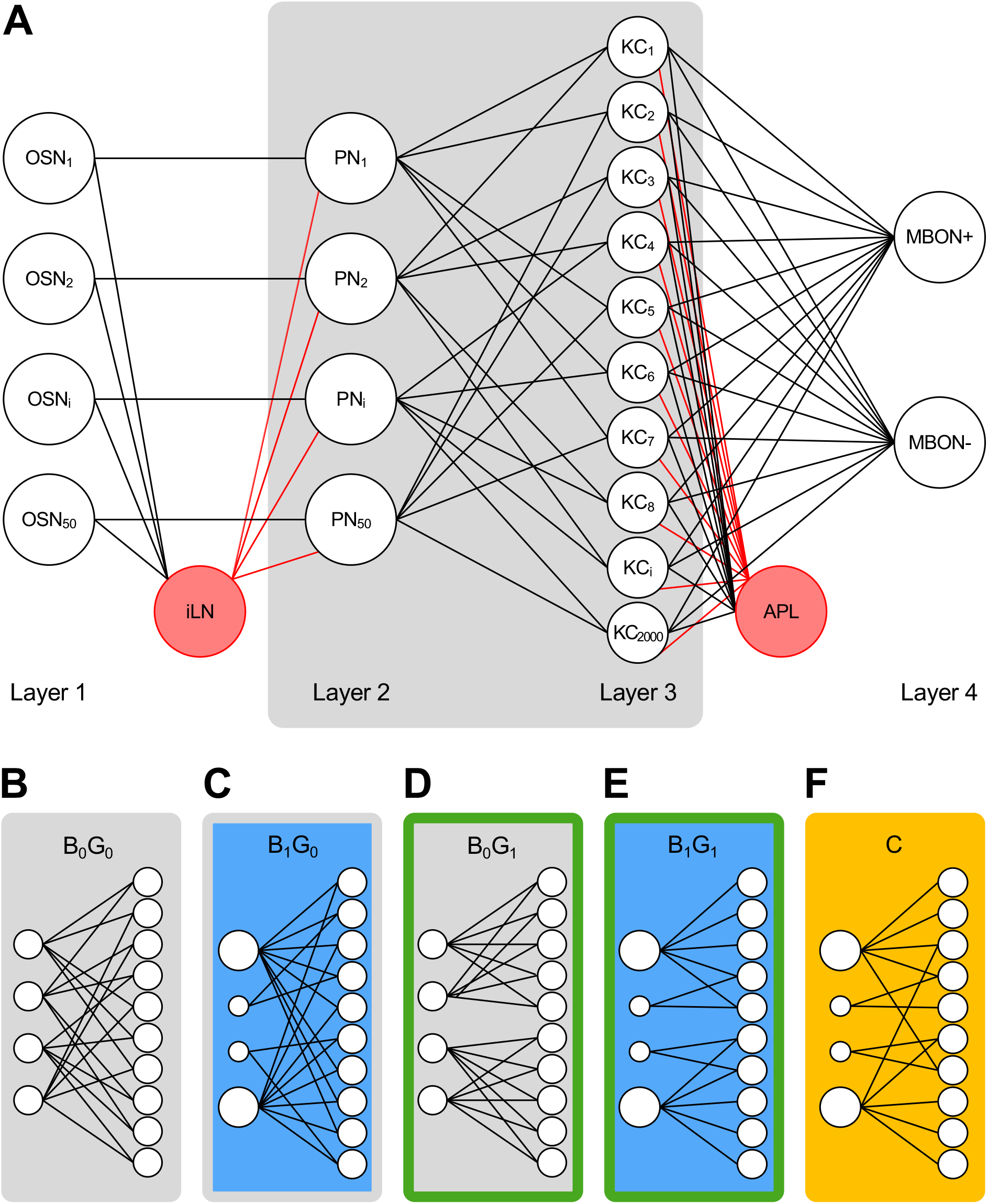
A four-layer network that replicates features of the *Drosophila melanogaster* mushroom body. (A) A four-layer network replicates key features of the *Drosophila melanogaster* mushroom body: the first two layers consist of 100 neurons (50 OSNs and 50 PNs) that connect in a one-to-one manner; the third layer consists of 2000 KCs that each receives a small number of PN inputs; the fourth layer consists of two MBONs that each receive input from all KCs and mediates either attraction (MBON+) or repulsion (MBON-); a feedforward inhibitory neuron (iLN) sums the signal elicited in all OSNs and inhibits individual PNs; a feedback inhibitory neuron (APL) sums the signal elicited in all KCs and inhibits individual KCs; excitatory and inhibitory connections are represented as black and red lines, respectively. Different variants of this network were built by adding structure to the PN-KC connections (grey box). (B-F) The connections between PNs and KCs were either unstructured (B, grey fill and outline) or structured using biases (C, blue fill and grey outline), groups (D, grey fill and green outline), a combination of both connectivity patterns (E, blue fill and green outline) or structured based on the most recent mushroom body connectome (F, yellow fill and outline).

In line with the predictions obtained from these theoretical studies, experimental studies of the *Drosophila melanogaster* mushroom body — one of the best characterized cerebellum-like structures — have shown that the mushroom body represents odors as sparse and distributed activity patterns (Honegger et al., 2011; Lin et al., 2014). Counter to these predictions however, there is ample experimental evidence showing that the connections between the input neurons of the mushroom body — the projection neurons — and its encoding neurons — the Kenyon cells are largely, but not completely random (Caron et al., 2013; Eichler et al., 2017; Gruntman and Turner, 2013; Li et al., 2020a; Murthy et al., 2008; Zheng et al., 2018, 2020). Two principal connectivity patterns have been identified by which these connections diverge from complete randomness. First, projection neurons connect with Kenyon cells at different likelihoods such that some projection neurons form more connections with Kenyon cells than others. The result is a non-uniform distribution of Kenyon cell input that differs significantly from the uniform distribution that would be expected under complete randomness (Caron et al., 2013). Second, several studies suggested that individual projection neurons can be divided into groups based on morphological features, and that Kenyon cells are more likely to integrate input from the projection neurons that belong to the same group (Gruntman and Turner, 2013; Lin et al., 2007; Zheng et al., 2020). Below, we term these two connectivity patterns ‘biases’ and ‘groups’, respectively. Both connectivity patterns have been confirmed in the most recent connectome of the *Drosophila melanogaster* brain, however, their functional consequences remain largely unknown(Li et al., 2020a; Scheffer et al., 2020; Zheng et al., 2020).

Even though *Drosophila melanogaster* affords researchers with an unmatched toolkit for analyzing and manipulating the function of individual neurons within a circuit, it is much more difficult to alter experimentally larger-scale aspects of brain architecture and assess the functional consequences. We therefore resorted to computational modeling to investigate the possible network functions enabled by biases and groups. We built a four-layer network that simulates odor representation in the mushroom body. We generated ten variants of this network by changing the degree of structure between the modeled projection neurons (PNs) and the modeled Kenyon cells (KCs) using biases, groups or a combination of both connectivity patterns. One of these variants replicates the input connections identified in the most recent mushroom body connectome (Li et al., 2020a). We compared odor representations generated by the KC layer across these ten variants qualitatively and quantitatively; we also assayed the ability of a network to discriminate and learn odors. We found that the representation space generated by structured networks is of lower dimensionality when compared to the representation space generated by unstructured networks and, consequently, structured networks show weaker performance in an associative learning task. We also found that the two connectivity patterns endow a network with specific functions. First, biased networks enable the robust representation of a subset of ethologically meaningful odors — enhancing the ability of these networks to form association with these odors — while disabling the representation of another subset of ethologically meaningful odors, rendering these odors unlearnable. Second, grouped networks have a greater capacity to generalize learning across similar odors while maintaining the ability to discriminate across most odors. Altogether, our study demonstrates that biases and groups are connectivity patterns that organize the representation space of the mushroom body, possibly fine-tuning its ability to represent and learn ethologically meaningful odors.

## RESULTS

### A four-layer network that simulates odor representation in the *Drosophila melanogaster* mushroom body

The Kenyon cells — the constituent neurons of the mushroom body — represent odors as sparse and distributed activity patterns (Honegger et al., 2011). During associative learning, the connections between the Kenyon cells activated by an odor and the mushroom body output neurons mediating learned behaviors are modified (Cohn et al., 2015; Hige et al., 2015; Owald et al., 2015; Séjourne et al., 2011). We built a four-layer network replicating these functions of the mushroom body using different anatomical and functional features, as they have been defined experimentally in *Drosophila melanogaster* (Figure 1A). The Kenyon cells receive most of their inputs from the antennal lobe — the primary olfactory processing center divided into about 50 glomeruli — through the projection neurons (Marin et al., 2002; Wong et al., 2002). Individual glomeruli receive input from the olfactory sensory neurons that express the same receptor gene(s), and glomeruli are interconnected through a network of inhibitory local neurons (reviewed in Wilson, 2013). Reflecting this discrete organization of the antennal lobe, the first two layers of the network — which we termed ‘OSNs’ and ‘PNs’, respectively — each consists of 50 neurons connecting in a one-on-one manner. The model used to transform OSN activity patterns into PN activity patterns replicates functional features of the projection neurons (Bhandawat et al., 2007; Olsen and Wilson, 2008; Olsen et al., 2010; Wilson et al., 2004). As a minimal model of the global lateral inhibition occurring in the antennal lobe, we implemented an inhibitory local neuron — which we termed ‘iLN’ — that sums the activity of OSNs and inhibit all PNs; each PN differs in its sensitivity to inhibition, reflecting experimental data (Hong and Wilson, 2015).

The third layer of the network consists of 2,000 neurons — which we termed ‘KCs’ — and each KC receives input from a small number of PNs, as is the case in *Drosophila melanogaster* (Caron et al., 2013; Li et al., 2020a). The model used to transform PN activity patterns into KC activity patterns is based on several experimental studies (Gruntman and Turner, 2013; Turner et al., 2008). We also included in the KC layer an inhibitory interneuron — which we termed APL — that mediates all-to-all feedback inhibition between KCs. This neuron replicates features of the anterior paired lateral neuron and its homologous locust neuron, the giant GABAergic neuron (Lin et al., 2014; Papadopoulou et al., 2011). The fourth layer of the network consists of two neurons, ‘MBON+’ and ‘MBON-’, and each MBON sums input across all KCs. The neurons receiving the strongest input directs the behavior. These two MBONs are a simplification of the 35 mushroom body output neurons found in the *Drosophila melanogaster* brain that encode different learned behaviors, namely approach and avoidance (Aso et al., 2014a, 2014b).

We generated a total of ten variants of this four-layer network. Each variant differs in the way the PN layer connects to the KC layer (Figure 1B-F; Supplementary Figure 1). In nine of these variants, we structured the PN–KC connections using biases, groups or varying degrees of both connectivity patterns while keeping the average total number of PN–KC connections constant. Each variant is named based on the amount of bias and group structure it was built with (biases: B_0/0.5/1_; groups: G_0/0.5/1_). In the B_0_G_0_ variant, the PN–KC connections are completely unstructured and, therefore, distributed randomly and uniformly (Figure 1B). The B_0.5_ and B_1_ variants were built using biases. In these networks, each PN forms a different number of connections to KCs based on the distribution of Kenyon cell inputs that has been measured experimentally (Figure 1C, Supplementary Figure 1; Caron et al., 2013). In the B_0.5_ variant, the number of PN–KC connections formed by a given PN lies halfway between its values in the uniform and non-uniform distributions. The G_0.5_ and G_1_ variants were built using groups. In these networks, the PNs were divided into five groups. These groups were partially defined based on different anatomical studies (Figure 1D, Supplementary Figure 1; Jefferis et al., 2007; Lin et al., 2007). In the G_1_ variant, individual KCs receive input only from the PNs that belong to the same group; the number of KCs receiving input from a given group of PNs — 400 KCs — was kept constant across all five groups. In the G_0.5_ variant, only half of the KCs — 1000 KCs — were structured this way. In the tenth variant or C variant, we structured the PN–KC connections based on the most recent mushroom body connectome (Figure 1F; Li et al., 2020a). Thus, the PN–KC connections in a given network variant are either unstructured (B_0_G_0_; Figure 1B), bias-structured (B_0.5-1_; Figure 1C), group-structured (G_0.5-1_; Figure 1D), or structured using both connectivity patterns (B_0.5-1_G_0.5-1_ and C; Figure 1E,F). For each experiment, we generated *n* realizations of each network variant and reported the average result obtained for all the realizations of a given variant.

We used these network variants to determine whether and how the different types of connectivity pattern affect the way the mushroom body represents odors and forms associations. To this end, we simulated odor responses in the OSN layer using two different odor panels. The first panel — which we refer to as the ‘realistic odor panel’ — was built by assigning an activity pattern to each of the 50 OSNs based on the odor-evoked responses that have been recorded experimentally in olfactory sensory neurons (Hallem and Carlson, 2006; Münch and Galizia, 2016). The realistic odor panel contains simulations for 110 odorant molecules. Most of these odorant molecules were found to activate multiple types of olfactory sensory neuron; we named these ‘multi-odors’. We added to the realistic panel six odors simulating molecules that have been found in separate studies to activate only one type of olfactory sensory neuron; we named these ‘mono-odors’ (Dweck et al., 2013; Kurtovic et al., 2007; Ronderos et al., 2014; Schlief and Wilson, 2007; Stensmyr et al., 2012; Suh et al., 2004). These six odorant molecules — carbon dioxide, geosmin, pyrrolidine, *cis-*vaccenyl acetate, farnesol and valencene — play a role in a variety of innate and learned behaviors. The second odor panel — which we refer to as the ‘expanded odor panel’ — was built by arbitrarily assigning an activity pattern to each of the 50 OSNs; this odor panel is meant to resemble the realistic odor panel but to increase the total number of odors from 116 to 5,000.

### Structure reduces the dimensionality of the representation space of a network

As a first step to compare the different network variants, we measured the dimensionality of their representation space at the level of the KC layer. The dimensionality of the representation space is an estimate of the minimum number of coordinates needed to describe the activity patterns generated by a given network in response to a large set of stimuli, ideally all possible stimuli. High dimensionality indicates high linear separability — and therefore high discriminability between stimuli — whereas low dimensionality indicates low separability and discriminability between stimuli (Cayco-Gajic and Silver, 2019; Fusi et al., 2016; Huerta et al., 2004). We compared the dimensionality of the network variants using 2,000 odors from the expanded odor panel (Figure 2A). The highest dimensionality was obtained for the unstructured variant (B_0_G_0_: 0.85 ± 0.05, *n*=20; Figure 2A). Structure — regardless of the type — consistently reduces the dimensionality of the representation space. However, the variants built using biases maintain a relatively high dimensionality, compared to other structured variants, suggesting that biases have a minor effect on the overall ability of a network to generate a large number of separable representations (B_0.5_G_0_: 0.80 ± 0.06, *n*=20; B_1_G_0_: 0.65 ± 0.04, *n*=20; Figure 2A). The variant built based on the mushroom body connectome also maintains high dimensionality (C: 0.68 ± 0.05, *n*=20; Figure 2A). In contrast, the variants built using groups show a drastic reduction in their dimensionality, suggesting that these networks can generate far fewer separable representations (B_0_G_0.5_: 0.55 ± 0.10, *n*=20; B_0_G_1_: 0.33 ± 0.08, *n*=20; Figure 2A). In line with these findings, the lowest dimensionality score was obtained for the most structured variant (B_1_G_1_: 0.24 ± 0.04, *n*=20; Figure 2A).

**Figure 2.**
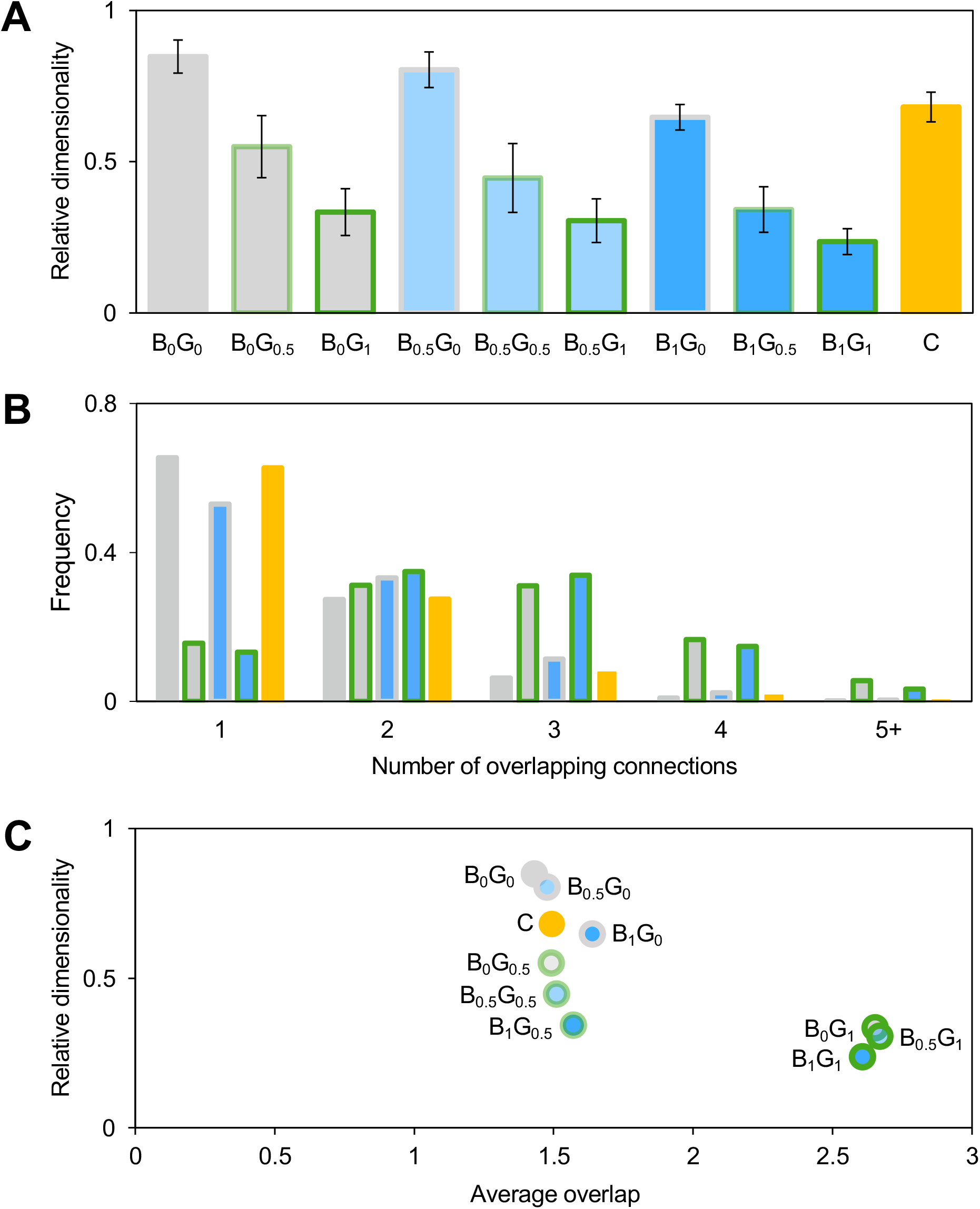
Structure reduces dimensionality and increases input overlap. (A) Each of the ten network variants were used to generate 2,000 KC representations using the expanded odor panel; the relative dimensionality of the resulting 2,000 representations was measured and averaged across network realizations (*n*=20). Error bars represent the standard deviation. (B) The frequency of overlapping connections among KCs — as defined by KC pairs that share one (1), two (2), three (3), four (4), five or more (5+) common PN inputs — was aggregated across network realizations (*n*=10). Note that the probability of having zero overlapping inputs is not shown in the distribution. (C) The average relative dimensionality obtained for each network variant is represented as a function of the average frequency of overlapping connections obtained for that variant (*n*=10). Grey fill and outline: unstructured variant; blue fill: variants built using biases (light blue fill: B_0.5_; dark blue fill: B_1_); green outline: variants built using groups (light green outline: G_0.5_; dark green outline: G_1_); yellow fill and outline: variant built based on the mushroom body connectome.

It is conceivable that the lower dimensionality observed with the structured variants results from an increase in the number of KCs that receive the same complement of inputs in these networks. Indeed, a previous study found that the dimensionality of cerebellum-like networks varies with the amount of input overlap among encoding neurons: the more overlap there is, the lower will be the dimensionality of a network (Litwin-Kumar et al., 2017). We thus tested whether the dimensionality of the different network variants is inversely proportional to the amount of input overlap among KCs. To this end, we measured the frequency of overlapping connections — the probability of two KCs sharing two, three, four or five and more inputs — in a given network (Figure 2B-C). We found that, for the B_0_G_0_, B_1_G_0_ and C variants, the frequency of overlapping inputs decreases monotonically (Figure 2B). The average input overlap across KC pairs is only slightly higher in the B_1_G_0_ and C variants than it is in the unstructured variant (B_0_G_0_: 1.4, *n*=10; B_1_G_0_: 1.6, *n*=10; C: 1.5, *n*=10; Figure 2C). In contrast, the frequency of overlapping inputs does not decrease monotonically in the B_0_G_1_ and B_1_G_1_ variants but, instead, reaches a maximum value at two and three shared inputs (Figure 2B). The average input overlap across KC pairs in these variants is much higher (B_0_G_1_: 2.7, *n*=10; B_1_G_1_: 2.6, *n*=10; Figure 2C). This observation is in line with our finding that the B_0_G_0_, B_0.5_G_0_, B_1_G_0_ and C variants generate a representation space of higher dimensionality, implying a negative correlation between the dimensionality of the representation space and the average amount of input overlap. However, we observed that the variants in which only half of the KCs are group-structured — specifically the B_0_G_0.5_, B_0.5_G_0.5_ and B_1_G_0.5_ variants — behave differently. Although these network variants have a relatively low dimensionality, their average input overlap is comparable to that of the completely unstructured variant (B_0_G_0.5_: 1.5, *n*=10; B_0.5_G_0.5_: 1.5, *n*=10; B_1_G_0.5_: 1.6, *n*=10; Figure 2C). This observation suggests that — although the reduction in dimensionality observed in the G_1_ variants correlates with an increased average overlap in KC inputs — other network features contribute to lowering dimensionality.

Likewise, it is conceivable that the lower dimensionality observed with the structured variants results from differences in the density of the representations generated by these networks (see next section; Figure 5A). Indeed, a previous study demonstrated that the dimensionality of cerebellum-like networks is related to the density of the representations they generate: the sparser representations are, the higher will be the dimensionality (Litwin-Kumar et al., 2017). We thus tested whether the trend in dimensionality observed for the different network variants is maintained when the density of odor representations — the % of KCs activated by an odor — is fixed at a given value. To this end, we measured the relative dimensionality of the representation space generated by each network variant while fixing the percentage of KCs activated at an odor at values ranging from 1% to 50% of the KCs. We found that the trend we detected across variants is observed across all density values (Supplementary Figure 2). Namely, the B_0_G_0_ variant has the highest dimensionality and the B_1_G_1_ variant has the lowest dimensionality; the effect of groups on dimensionality is greater than the effect of biases. These trends are especially prominent at realistic density values when the density is fixed at 5, 10 or 15 % (Supplementary Figure 2). These observations suggest that the effect of structure on dimensionality is not strictly caused by differences in the density of the representations generated by a network.

Altogether, these results show that structure — either in the form of biases or groups — reduces dimensionality; the reduction in dimensionality observed in structured networks is, in part, caused by an increase in the number of overlapping inputs among KCs but not strictly caused by differences in the density of the representations generated by these networks.

### Structure affects the recall capacity of a network

Ultimately, the function of the mushroom body is to enable a fly to learn. It does so by forming associations between odors and different behavioral outcomes and by recalling these associations. It is conceivable that the dimensionality of the representation space generated by a network affects its learning performance: networks of higher dimensionality could, in principle, enable better learning performance and *vice versa*. We tested this idea by devising a learning task based on previous theoretical and experimental studies (Peng and Chittka 2017; Cohn et al. 2015; Handler et al. 2019). This learning task consists of two steps: a training step and a testing step (Figure 3A). During the training step, the network forms a number of associations; for each association, a behavioral outcome — approach or avoidance — is randomly assigned to an odor selected from the expanded odor panel. Learning was implemented in the network by modifying the weight between KCs and MBONs, such that it reflects currently available experimental data (Cohn et al., 2015; Handler et al. 2019). Specifically, the connections between the KCs activated by an odor and the MBON mediating the opposite behavior were weakened by an *α*_LTD_ value, whereas the connections between the non-active KCs and that same MBON were strengthened by an *α*_LTP_ value; the connections between the activated KCs and the MBON mediating the learned behavior were left unchanged. During the testing step, the trained network was presented with the odors it had previously learned: a recall was considered correct when the behavioral response of the network to an odor was the one it had previously learned; failure to do so was considered an incorrect recall. We repeated the training step for *x* odors (*x*=20, 40, 60, …, 1200) and tested how many of the *x* associations a network had formed could be recalled correctly. ‘Recall accuracy’ represents the ratio of associations a given network could recall correctly: a recall accuracy of 1 indicates that the network performed perfectly and could recall all the associations it had formed whereas a recall accuracy of 0.5 indicates that that the performance of the network was just as good as that of an untrained network.

**Figure 3.**
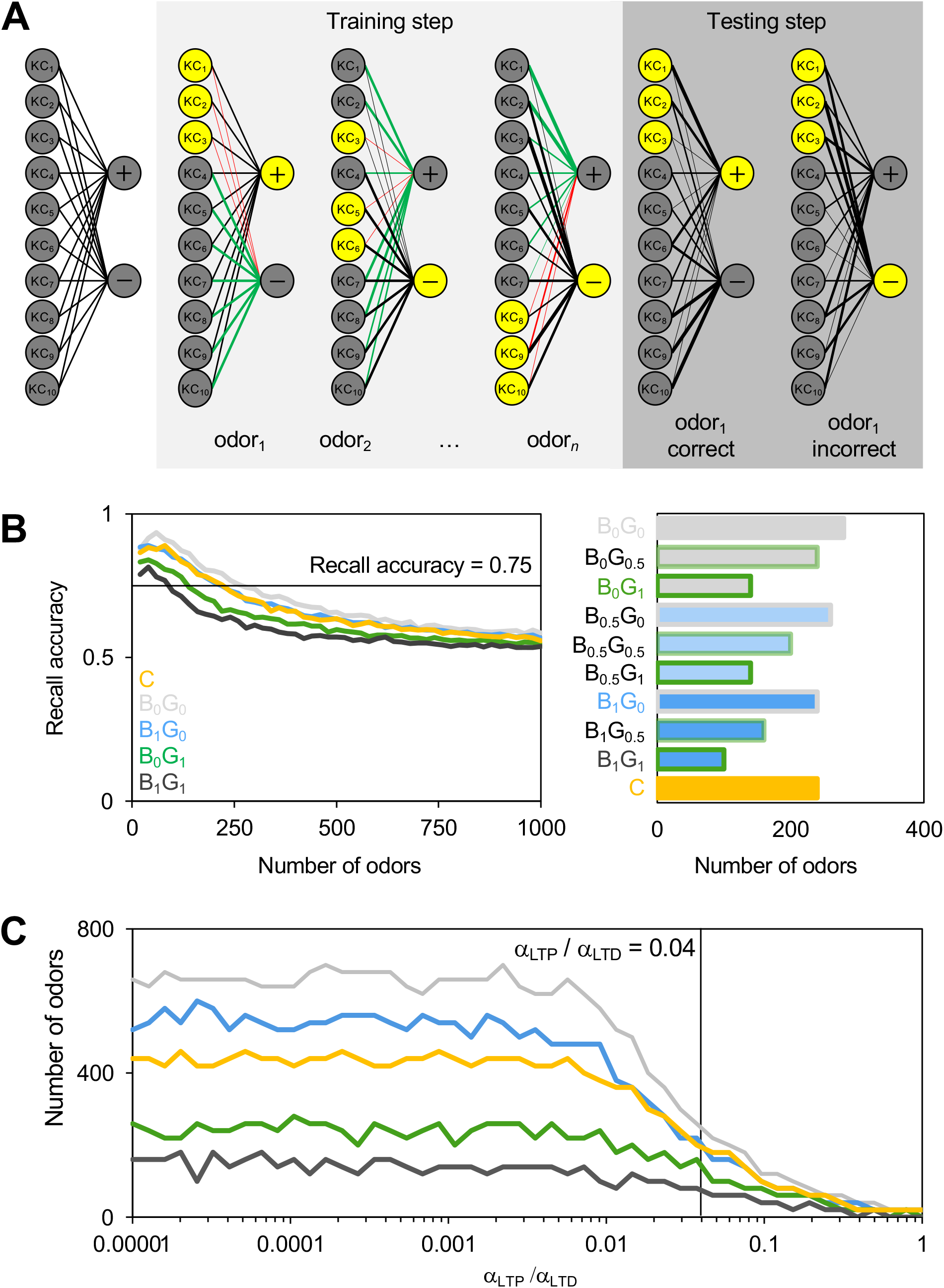
Structure affects recall accuracy. (A) Each MBON receives input from all KCs and mediates either attraction (+) or repulsion (-). During the training step (light grey box), at the presentation of each odor, the weight of the connections between the KCs activated by this odor (yellow KCs) and the MBON driving the opposite behavior (grey MBON) are reduced by a *α*_LTD_ value (red lines), whereas the weight of the connections between inactive KCs (grey KCs) and the same MBON (grey MBON) is increased by an *α*_LTP_ value (green lines). During the testing step (dark grey box), the trained network is presented with an odor it had previously learned; a correct recall means that the MBON activated by the odor is the same as was associated with this odor in the training step, whereas an incorrect recall means that the other MBON was activated. (B) Left panel: the fraction of odors recalled correctly by a network, or ’recall accuracy’, as a function of the number of odors the network was trained with (*n*=20). Light grey line: B_0_G_0_ variant; blue line: B_1_G_0_ variant; green line: B_0_G_1_ variant; dark grey line: B_1_G_1_ variant; yellow line: C variant. Right panel: the number of odors the different network variants were trained with such that their recall accuracy was 0.75 is shown (intersection of the black line in the left panel). Grey fill and outline: unstructured variant; blue fill: variants built using biases (light blue fill: B_0.5_; dark blue fill: B_1_); green outline: variants built using groups (light green outline: G_0.5_; dark green outline: G_1_); yellow fill and outline: variant built based on the mushroom body connectome. (C) The number of odors the different network variants were trained with such that their recall accuracy was 0.75 as a function of the ratio of *α*_LTP_ to *α*_LTD_ is shown (*n*=5). Light grey line: B_0_G_0_ variant; blue line: B_1_G_0_ variant; green line: B_0_G_1_ variant; dark grey line: B_1_G_1_ variant; yellow line: C variant. The vertical black line indicates the values obtained when *α*_LTP_/*α*_LTD_ is 0.04 as in (B).

We compared the recall accuracy of different network variants by increasing the number of odors used during the training step (Figure 3B, left panel), as well as by quantifying the number of odors each of the variants could recall, such that its recall accuracy was 0.75 (Figure 3B, right panel). The B_0_G_0_ variant performed best — its recall accuracy falls below 0.75 after forming associations with 280 odors — indicating that, as predicted, unstructured networks can achieve the greatest recall capacity. The B_1_G_0_ variant performed nearly as well: its recall accuracy falls below 0.75 after forming 240 associations indicating that biases have minimal impact on the performance of a network. The C variant performed in a similar fashion and its recall accuracy falls below 0.75 after forming 240 associations. In contrast, we observed a more notable decline in performance with the B_0_G_1_ variant: its recall accuracy falls below 0.75 after forming 140 associations, indicating that group-structure has a substantial impact on the recall capacity of a network. The weakest performance was achieved by the most structured variant, B_1_G_1_: its recall accuracy falls below 0.75 after forming only 100 associations. We investigated whether the strength of the *α*_LTD_ to *α*_LTP_ ratio influences the performance of a network by repeating the analysis described above while holding the *α*_LTD_ value constant and varying the *α*_LTP_ value (Figure 3C). Indeed, lower *α*_LTP_/*α*_LDP_ ratios enable better learning performance as the recall capacity of a network increases monotonically when the *α*_LTP_/*α*_LDP_ ratio decreases, and this across network variants; the recall capacity eventually plateaus when *α*_LDP_ is three orders of magnitude stronger than *α*_LTP_. Our initial observation — whereby biases affect learning performance but less so than groups — is consistent across all *α*_LTP_/*α*_LDP_ ratios.

Altogether, these results demonstrate that the learning performance of a network in a recall task correlates with the dimensionality of its representation space: networks generating representations of high dimensionality perform better than networks generating representations of low dimensionality.

### Structure enables the representation of mono-odors

Although, unstructured networks undoubtedly perform best in a recall task, they do not accurately represent the *Drosophila melanogaster* mushroom body: several anatomical studies found that the connections between projection neurons and Kenyon cells are largely, but not completely, random (Caron et al., 2013; Gruntman and Turner, 2013; Li et al., 2020; Zheng et al., 2018, 2020). To identify the possible functions enabled by these connectivity patterns, we investigated the performance of the network variants in more realistic scenarios. We analyzed the representations generated by the different variants using the realistic odor panel (Figure 4A). We found that the percentage of KCs activated by multi-odors — odors that activate several OSNs — slightly differs across variants: the unstructured variant generates relatively sparse representations (B_0_G_0_: 8.2% ± 1.8 of the KCs, *n*=50; Figure 4A) whereas the most structured variant generates slightly denser representations (B_1_G_1_: 13.7% ± 4.3 of the KCs, *n*=50; Figure 4A). Each connectivity pattern has a different effect on the density of representations: increasing levels of group-structure leads to denser representations (B_0_G_0.5_: 10.0% ± 2.7 of the KCs*, n*=50; B_0_G_1_: 12.4% ± 3.5 of the KCs, *n*=50; Figure 4A), whereas increasing the level of bias-structure leads to sparser representations (B_0.5_G_0_: 7.6% ± 2.1 of the KCs, *n*=50; B_1_G_0_: 7.5% ± 2.0 of the KCs, *n*=50; Figure 4A). The C variant generates representations that are comparable in density to the ones generated by the biased variants (C: 6.3% ± 1.6 of the KCs, *n*=50; Figure 4A). Despite these subtle but noticeable differences, the values obtained for all network variants are consistent with the values obtained experimentally, as most odors activate on average between 5 and 15% of the Kenyon cells (Honegger et al., 2011; Lin et al., 2014).

**Figure 4.**
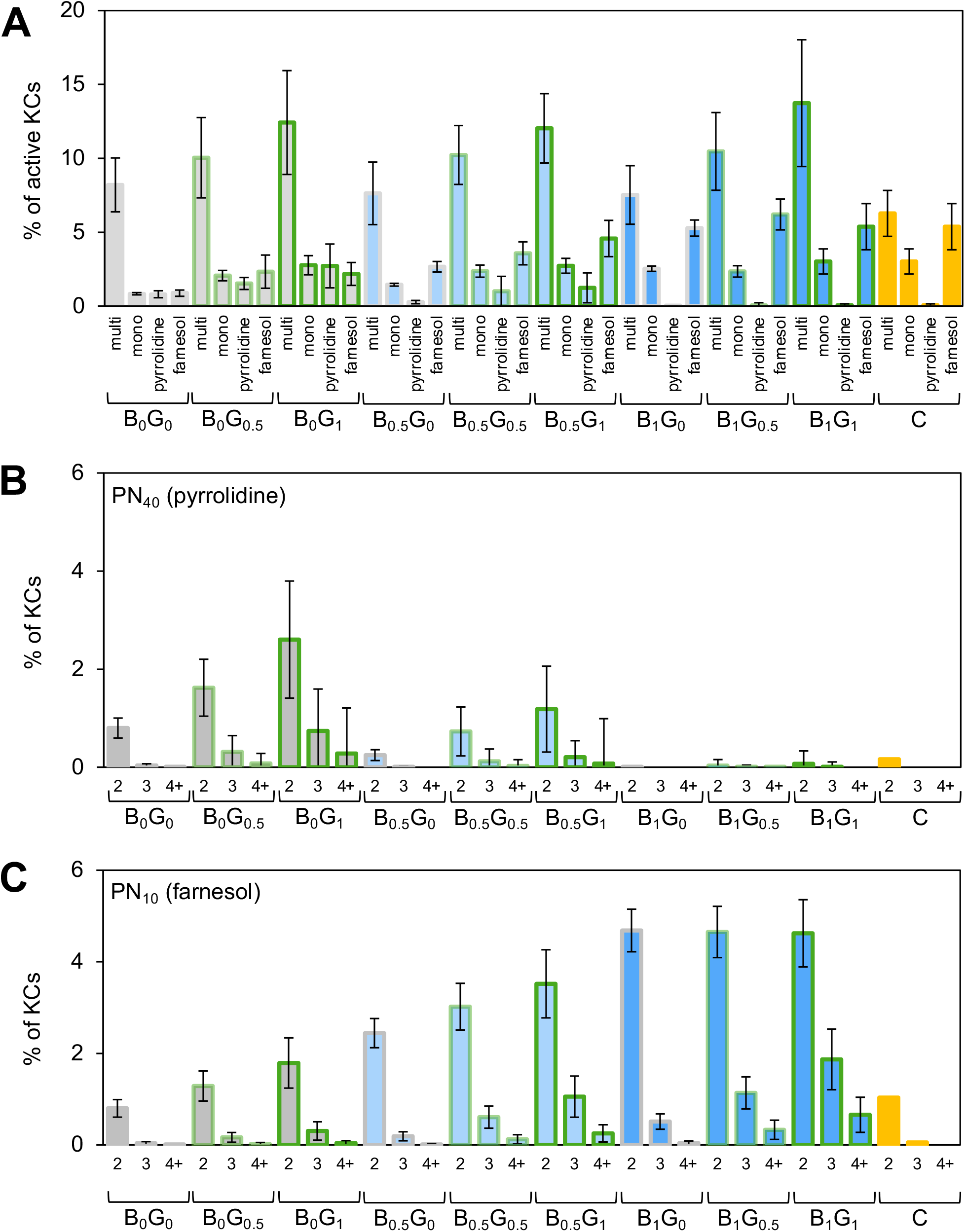
Structure enables the representation of mono-odors. (A) The percentage of KCs activated by different types of odor — odors activating either multiple OSNs (multi), one OSN (mono), one OSN that connects to PN_40_, which is underrepresented in biased networks (pyrrolidine) or one OSN that connects to PN_10_, which is overrepresented in biased networks (farnesol) — was measured in each of the ten variants and averaged (*n*=50). (B) The percentage of KCs that receive two (2), three (3), four and more (4+) inputs from PN_40_ — which is activated by pyrrolidine — was measured across all ten variants and averaged (*n*=500). (C) The percentage of KCs that receive two (2), three (3), four and more inputs (4+) from PN_10_ — which is activated by farnesol — was measured across all ten variants and averaged (*n*=500). Error bars represent the standard deviation. Grey fill and outline: unstructured variant; blue fill: variants built using biases (light blue fill: B_0.5_; dark blue fill: B_1_); green outline: variants built using groups (light green outline: G_0.5_; dark green outline: G_1_); yellow fill and outline: variant built based on the mushroom body connectome.

In contrast, we found that the percentage of KCs activated by mono-odors — odors that activate only one OSN — varies considerably across variants. Mono-odors generate extremely sparse representations in the B_0_G_0_ variant, a value below the values obtained for multi-odors (B_0_G_0_: 0.8% ± 0.1 of the KCs, *n*=50; Figure 4A). Both connectivity patterns enable slightly denser representations for mono-odors but these representations are significantly sparser than those obtained for multi-odors (B_1_G_0_: 2.5% ± 0.2 of the KCs, *n*=50; B_0_G_1_: 2.8% ± 0.6 of the KCs, *n*=50; C: 3.0% ± 0.9 of the KCs, *n*=50; Figure 4A). Interestingly, the PNs activated by mono-odors are among those that form the most or the fewest connections to KCs in the B_1_G_0_ and C variant. This observation raised the possibility that these variants might represent mono-odors differently. To investigate this possibility further, we analyzed in more detail the representations formed by two mono-odors: pyrrolidine — an odor that activates PN_40_, which forms fewer connections than average with KCs in the B_1_G_0_ and C variants — and farnesol — an odor that activates PN_10_, which forms more connections than average with KCs in these variants. We observed that these odors generated completely different representations: in the B_1_G_0_ and C variants, pyrrolidine fails to activate any KCs, whereas farnesol generates representations that are nearly as dense as the ones generated for multi-odors (B_1_G_0_: pyrrolidine: 0.004% ± 0.014 of the KCs, *n*=50, farnesol: 5.3% ± 0.55 of the KCs, *n*=50; C: pyrrolidine: 0.06% ± 0.09 of the KCs, *n*=50, farnesol: 5.4% ± 1.6 of the KCs, *n*=50; Figure 4A). Thus, groups and biases enable the representation of mono- odors in different ways: groups enable the representations of all mono-odors but these representations are considerably sparser than those generated by multi-odors; biases enable the representations of some odors such as farnesol — with representations that are as dense as those generated for multi-odors — while disabling the representation of other odors such as pyrrolidine.

All network variants were tuned such that two or more active inputs are required to activate a KC, as has been determined experimentally for Kenyon cells (Gruntman and Turner, 2013). Thus, the density of the representation generated for a mono-odor most likely depends on the number of KCs that receive two or more inputs from the PN activated by the mono-odor. To verify this, we measured the number of homotypic connections — as defined by a PN connecting twice or more often to the same KC — in the different network variants (Figure 4B,C). We focused our analysis on the homotypic connections formed with the PN activated by pyrrolidine, PN_40_, as well as the PN activated by farnesol, PN_10_. Indeed, we found that such double homotypic connections with either PN_10_ or PN_40_ are rare in the unstructured variant (B_0_G_0_: PN_10_: 6.1% ± 1.5 of the KCs, *n*=500, PN_40_: 6.1% ± 1.5 of the KCs, *n*=500; Figure 4B,C). As expected, we found that the B_1_G_0_ and C variants show more double homotypic connections with PN_10_ than with PN_40_ (B_1_G_0_: PN_10_: 14.8% ± 1.5 of the KCs, *n*=500, PN_40_: 0.6 ± 1.6 of the KCs, *n*=500; C: PN_10_: 1.0% of the KCs, PN_40_: 0.16% of the KCs; Figure 4B,C). It is worth noting that triple, quadruple and quintuple homotypic connections are extremely rare, but less so in group-structured networks (Figure 4B,C). Thus, both connectivity patterns increase the frequency of homotypic connections; whereas biases primarily affect the number of double homotypic connections formed with a few PNs, groups increase the number of double, triple and quadruple connections formed with all types of PN.

Altogether, these results show that structure — regardless of its type — affects the representations of mono-odors specifically: biases enable the representation of a subset of mono- odors and disable the representation of different subset of mono-odors whereas groups enable the representation of all mono-odors.

### Biases prioritize the robust representation of a few mono-odors

Next, we investigated whether the representations generated by structured variants for mono- odors differ from those they generate for multi-odors. In particular, we wanted to test the robustness of these representations in the presence of noise. It is conceivable that the representations of mono-odors — because they depend on a single OSN — are more susceptible to noise. Alternatively, because the OSNs detecting these mono-odors are narrowly tuned to one or a few odors — and thus dedicated to the detection of one particular stimulus — it is possible that the representations generated by mono-odors are more robust. To simulate realistic, noisy input conditions, we added different intensities of Gaussian noise to the input in the OSN layer. We did so for twelve different odors selected from the realistic odor panel: two mono-odor, pyrrolidine and farnesol, as well as ten randomly selected multi-odors. We simulated representations for these twelve odors using the different network variants and ten realizations of noise for each odor. To quantify how robust the representations generated by a network are, we applied the nonlinear dimension reduction algorithm UMAP in combination with a k-means clustering tool (Figure 5A, left panels). UMAP was used to reduce the dimension of the representations from 2,000 to 2 while preserving both local and global relationships between data points (McInnes et al., 2018). The k-means clustering tool was used to divide the UMAP two-dimensional representations into 12 clusters, one for each odor.

**Figure 5.**
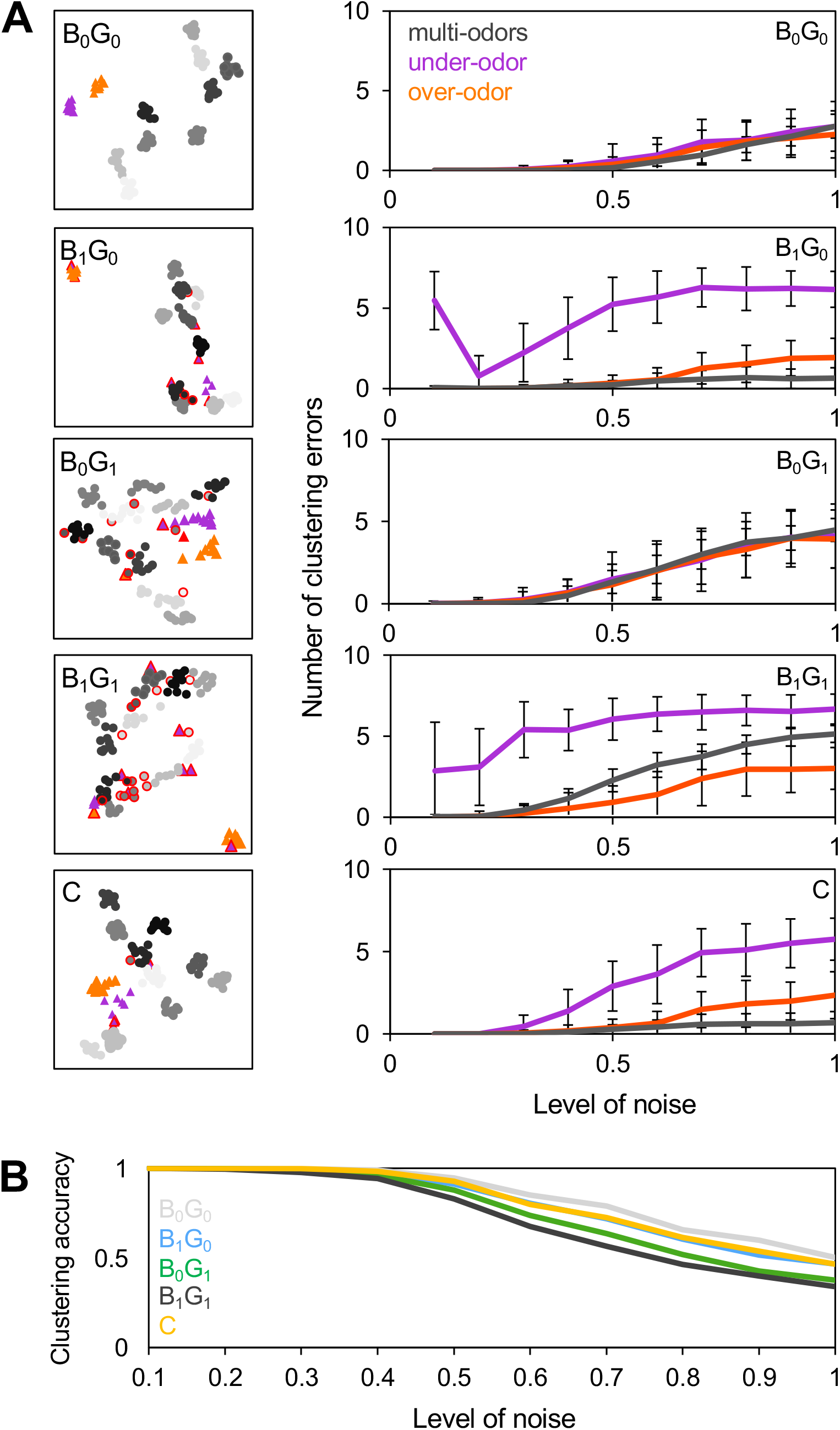
The representations of some mono-odors generated by biased-structured networks are robust to noise. (A) Left panels: KC representations were generated for 12 odors — farnesol (orange triangles), pyrrolidine (purple triangles) and ten multi-odors (grey circles, a given shade of gray represents one odor) — across network variants at a noise level of 0.5. Using the UMAP algorithm, the dimensionality of the representations generated for a given network was reduced from 2,000 to two; the resulting two UMAP dimensions serves as coordinates in the plot. The UMAP plots were analyzed using a clustering tool; the representations for a given odor that failed to cluster together are labeled (red outline). Right panels: The number of clustering incidents obtained for three different odors (purple: pyrrolidine, orange: farnesol, grey: a multi-odor) as a function of the level of noise is shown (*n*=50). (B) The average clustering accuracy (as measured for all 12 odors) as a function of the noise level is shown (*n*=50). Light grey line: B_0_G_0_ variant; blue line: B_1_G_0_ variant; green line: B_0_G_1_ variant; dark grey line: B_1_G_1_ variant; yellow line: C variant.

We then quantified the robustness of the representations generated by each network variant using two measurements. First, we measured the Rand index or ‘clustering accuracy’. This value reflects the success rate of the clustering tool in classifying the representations generated by the same odor as one cluster (Figure 5B). Second, we measured the number of mistakes made by the clustering tool for a given odor (Figure 5A, right panels). As expected, at low noise levels, all network variants produced representations that clustered well, but at high noise levels the quality of the clustering decreases significantly (Figure 5B). The unstructured B_0_G_0_ variant performs best, keeping higher clustering accuracy even at high noise levels and the most structured B_1_G_1_ variant performs worst. In line with our observations on the dimensionality of the representation space, we found that variants built using groups perform worse than those built using biases: while the B_1_G_0_ variant performs nearly as well as the unstructured B_0_G_0_ variant, the clustering accuracy for the representations generated by the B_0_G_1_ variant falls rapidly when noise is increased. The C variant performed most similarly to the B_1_G_0_ variant, maintaining high clustering accuracy even at high noise levels. Interestingly, the representations generated by the B_1_G_0_ and C variants for farnesol are nearly as robust than the ones generated for multi-odors (Figure 5A, right panels, orange line). In contrast, the representations generated by these variants for pyrrolidine are most susceptible to noise (Figure 5A, right panels, purple line). This is because these representations contain very few KCs, in some cases none. Thus, these results suggest that structure, in particular biases, enables the robust representation of some mono-odors.

Finally, we tested whether the structured variants could recall associations formed with mono-odors better than the unstructured variant can (Figure 6). To this end, we determined whether the recall accuracy obtained for mono-odors differed across network variants. We trained each network using two mono-odors, pyrrolidine and farnesol, as well as a number of odors randomly selected from the expanded odor panel. The total number of odors used during the training step was such that the recall accuracy of the trained network was 0.75. Thus, the number of odors used during training varied across variants (B_0_G_0_: 280 odors; B_1_G_0_: 240 odors; B_0_G_1_: 140 odors; B_1_G_1_: 100 odors; C: 240 odors; Figure 3B, right panel). We measured the recall accuracy using in the testing step either the mono-odors or the multi-odors that appeared near the mono-odors in the training sequence. As expected, the average recall accuracy obtained for multi-odors is approximately 0.75 across variants. However, the variants performed differently when recalling mono-odors. The B_0_G_1_ variant recalled the two mono-odors slightly better than it recalled multi- odors (B_0_G_1_: pyrrolidine: 0.80 ± 0.41, *n*=20 and farnesol: 0.90 ± 0.31, *n*=20; Figure 6) whereas the B_0_G_0_ variant failed at recalling both mono-odors (B_0_G_0_: pyrrolidine: 0.60 ± 0.50, *n*=20 and farnesol: 0.40 ± 0.50, *n*=20; Figure 6). The B_1_G_0_ and the B_1_G_1_ variants performed the best at recalling farnesol: these variants recalled correctly almost all the associations they had formed with farnesol (B_1_G_0_: 0.95 ± 0.22, *n*=20; B_1_G_1_: 0.95 ± 0.22, *n*=20; Figure 6). In contrast, the B_1_G_0_ and B_1_G_1_ variants failed at recalling pyrrolidine (B_1_G_0_: 0.50 ± 0.16, *n*=20; B_1_G_1_: 0.55 ± 0.36, *n*=20; Figure 6). In this recall test, the C variant performed most similarly to the B_0_G_0_ and the B_1_G_1_ variants as it could recalled associations made with farnesol better than it could recall associations made with pyrrolidine (C: pyrrolidine: 0.55 ± 0.51, *n*=20 and farnesol: 0.70 ± 0.47, *n*=20; Figure 6).

**Figure 6.**
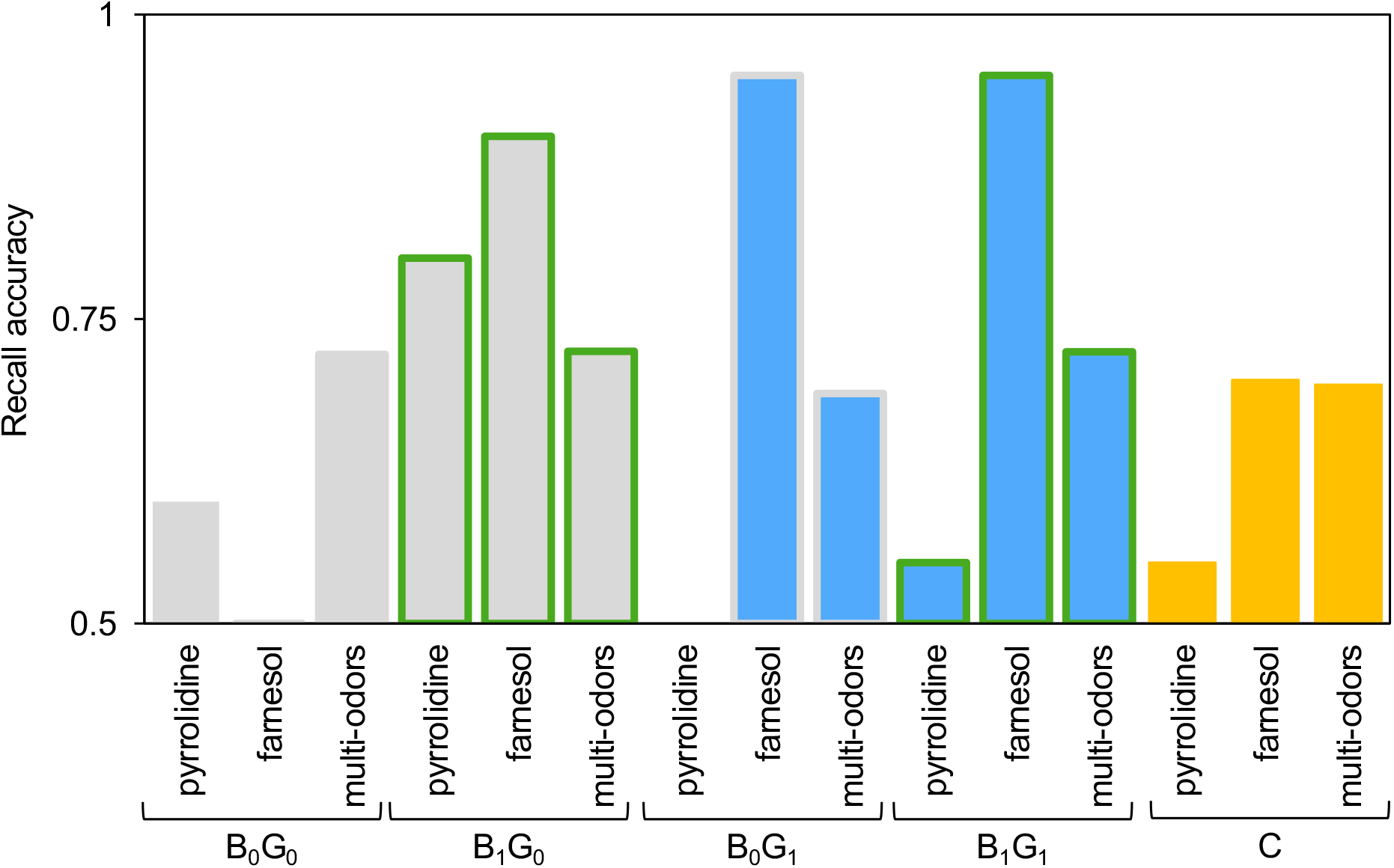
Biases-structured networks enable and disable the recall of some mono-odors. Different network variants were trained using pyrrolidine, farnesol and a number of multi-odors (randomly selected from the expanded odor panel) such that their recall accuracy was 0.75. The recall accuracy was measured separately for each odor (*n*=20). Grey fill and outline: unstructured networks; blue fill: networks built using biases (light blue fill: B_0.5_; dark blue fill: B_1_); green outline: networks built using groups (light green outline: G_0.5_; dark green outline: G_1_); yellow fill and outline: network built based on the connectome.

Combined, these observations suggest that networks built using biases represent mono- odors differently than the other networks do. The representations generated by the B_1_G_0_ network variant for some mono-odors, such as pyrrolidine, are less robust to noise than the ones generated for other odors; the B_1_G_0_ variant also failed at recalling associations made with pyrrolidine. In other words, the B_1_G_0_ variant cannot represent and learn pyrrolidine. In contrast, the representations generated by the B_1_G_0_ variant for farnesol are as robust to noise as those generated for multi-odors. Moreover, this variant can recall associations made with farnesol more accurately than it can recall associations made with multi-odors. Interestingly, farnesol as well as other mono-odors that activate a large number of KCs in the biased variant — such as *cis*- vaccenyl acetate and valencene — are molecules used in many complex, social behaviors that require learning and the mushroom body (Dweck et al., 2013; Kurtovic et al., 2007; Ronderos et al., 2014). By contrast, pyrrolidine as well as other mono-odors that fail to activate KCs in the biased variant — such as geosmin and carbon dioxide — are molecules that trigger strong, fixed behaviors that do not require learning and therefore do not require the mushroom body (Schlief and Wilson, 2007; Stensmyr et al., 2012; Suh et al., 2004). Thus, biases might prioritize the representation space of the mushroom body: the representations of odors that need to be learned in various contexts might occupy a larger space than the representations of odors that do not need to be learned (see Discussion).

### Groups enable networks to generalize learning across similar odors

It has been proposed that the higher the dimensionality of a network, to the better it can generate representations that are highly discriminable (Fusi et al., 2016). Because dimensionality varies greatly across network variants, it is possible that the ability of a network to generate highly discriminable representations varies too. To test the effect of structure on discriminability and generalizability, we performed pairwise comparisons of the representations generated by a given network variant. We evaluated the discriminability of a pair of representations using cosine distance as a measurement. A cosine distance of zero indicates that the two representations are identical, up to scalar multiplication, and therefore completely overlapping and hard to discriminate (Figure 7A, left panel). A cosine distance of one indicates that the two representations are orthogonal and therefore completely non-overlapping and highly discriminable (Figure 7A, right panel). Intermediate values of cosine distances indicate that the two representations overlap both in the identity of the KCs activated as well as their level of activation (Figure 7A, middle panel). We measured the cosine distance for all pairs of representations generated by a given network variant using 2,000 odors from the expanded odor panel (Figure 7B). The frequency of the cosine distances obtained using the B_0_G_0_ variant follows a unimodal distribution with a median value of 0.61. Surprisingly, the distribution obtained using the B_1_G_0_ variant is also unimodal but shifted towards higher values, with a median value of 0.67. This result indicates that — counter to the conjecture that dimensionality correlates with discriminability — networks built using biases have a greater ability to generate pairs of representations with higher cosine distances, and thus these networks can generate more representation pairs that have higher discriminability. The distribution obtained using the C variant resembles most the B_1_G_0_ distribution. The distribution obtained by comparing pairs of representations generated by the B_0_G_1_ variant is wider, with a lower median value of 0.55. The net result of this wider distribution is that this variant generates more pairs of representations with either very high cosine distances (higher than 0.92) or very low cosine distances (lower than 0.23). Such pairs are rarely found among the representations generated by the B_0_G_0_, B_1_G_0_ and C variants. This observation suggests that the B_0_G_1_ variant — despite having relatively low dimensionality — can generate representations with the widest range of discriminability, producing highly similar or highly dissimilar odor representations. Finally, the distribution of cosine distances obtained for the representations generated by the B_1_G_1_ variant has a median value of 0.61 and resembles the distribution obtained using the B_0_G_1_ variant, but with a noticeable shift towards higher values. Thus, the B_1_G_1_ and C variants combines features observed in both the B_1_G_0_ and B_0_G_1_ variants.

**Figure 7.**
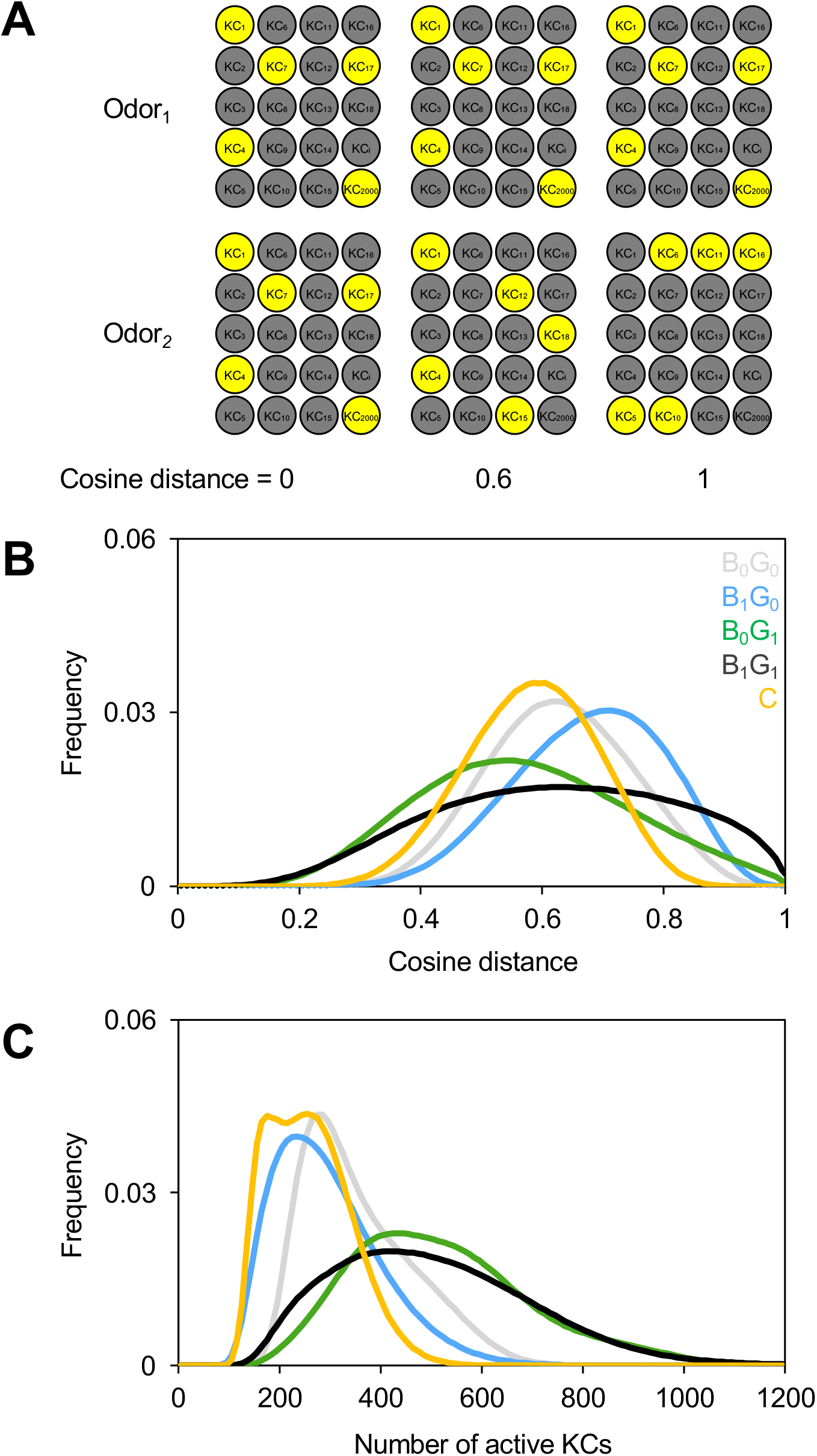
Group-structured networks generate a wider range of representation similarities. (A) Cosine distance measures the fraction of activated KCs (yellow KCs) shared between a pair of odor representations (upper panels: representations of odor_1_; lower panels: representations of odor_2_): a cosine distance of 0 means that there is complete overlap between the two representations (left panels) whereas a cosine distance of 1 means that there is no overlap between the two representations (right panels); intermediate values (here 0.6) reflect the number of KCs shared by the two representations (middle panels). (B) The distribution of cosine distances obtained for all pairs of representations generated by the different network variants is shown (*n*=20). Light grey line: B_0_G_0_ variant; blue line: B_1_G_0_ variant; green line: B_0_G_1_ variant; dark grey line: B_1_G_1_ variant; yellow line: C variant. (C) The distribution of the size of representations (number of activated KCs) generated by the different network variants is shown (*n*=20). Light grey line: B_0_G_0_ variant; blue line: B_1_G_0_ variant; green line: B_0_G_1_ variant; dark grey line: B_1_G_1_ variant; yellow line: C variant.

The wider range of representation similarities observed for group-structured networks most likely results from two different network features. First, because cosine distance directly measures the overlap between two representations, it must be related, in part, to the size of these representations. Odors that activate more KCs are more likely to generate overlapping representations. Indeed, the B_1_G_0_ variant has the highest median cosine distance (0.55) and generates sparse odor representations (multi-odors: 7.4% ± 2.0 of the KCs; Figure 5A and Figure 7C). In contrast, the B_0_G_1_ variant has the lowest median cosine distance (0.54) and generates denser odor representations (multi-odors: 12.4% ± 3 of the KCs; Figure 5A and Figure 7C). Second, group structure restricts the input of KCs to PNs that are assigned to the same group. Odors generating pairs of representations with very low cosine distances — representations that are greatly overlapping — must activate PNs that are within the same group, whereas odors generating pairs of representations with very high cosine distances — representations that show little overlap — must activate PNs distributed across different groups. To verify whether this is the case, we selected pairs of representations generated by the B_0_G_1_ variant in either the bottom or the top 2% of cosine distances (lower than 0.23 or above 0.92, respectively) and compared the PNs activated by each odor, in each pair. We found that the odors that generate representations of low cosine distance activate more PNs that belong to the same group than the odors that generate representations of high cosine distance (Figure 8A). It is worth noting that such pairs of representations — pairs with cosine distances lower than 0.23 or higher than 0.92 — cannot be found among the representations generated by the B_0_G_0_, B_1_G_0_ and C network variants. Because of these two network features — denser representations and restriction in KC inputs — group-structured networks are able to generate a wider range of representation similarities with, notably, a large fraction of highly similar representations.

**Figure 8.**
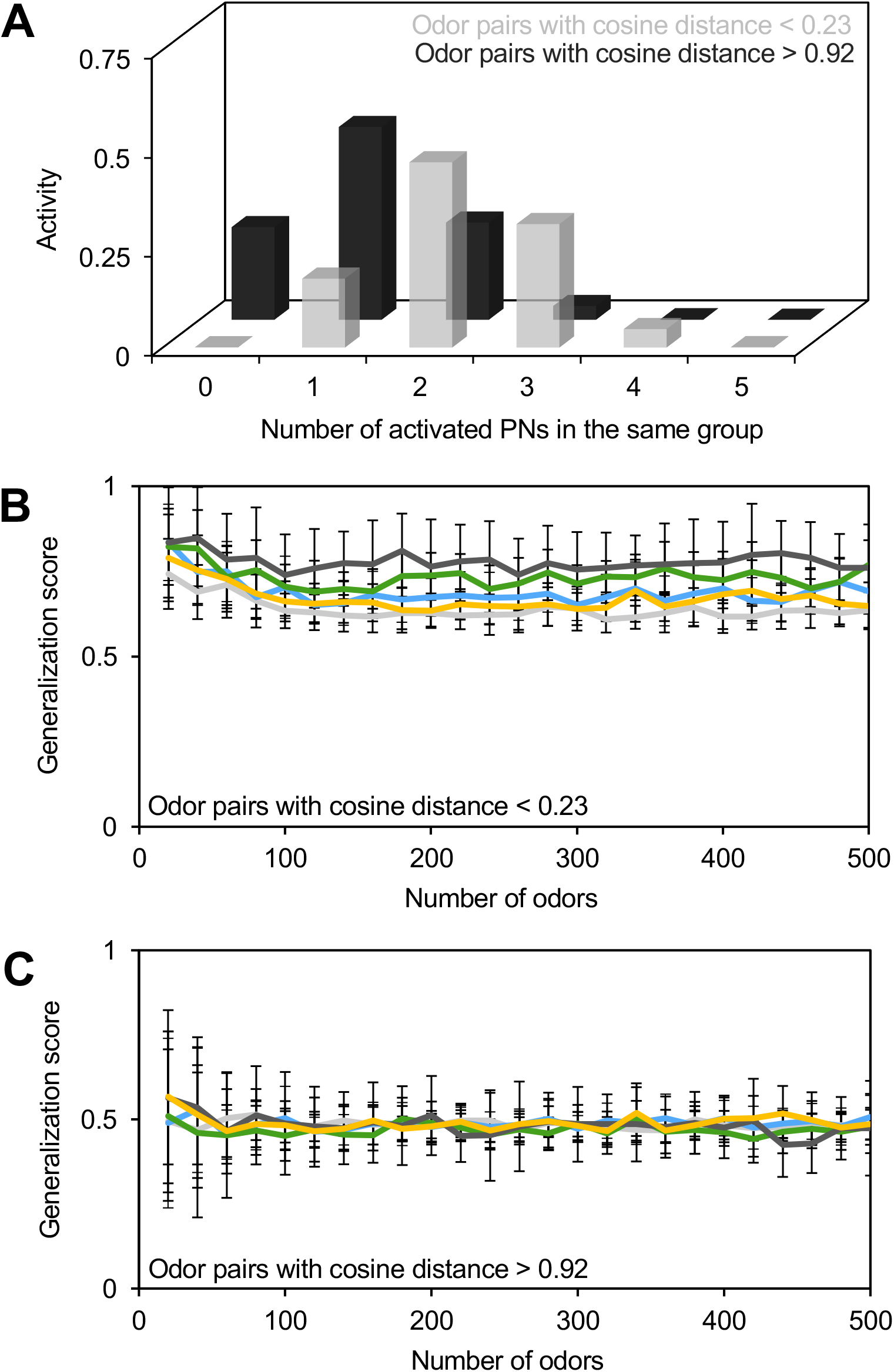
Group-structured networks can generalize across similar odors. (A) Pairs of odors generating representations with a low cosine distance (< 0.23) tend to activate PNs that belong to the same group (light gray) whereas pairs of odors generating representations with a high cosine distance (> 0.92) tend to activate PNs that belong to different groups (dark gray). (B-C) The ability of a network to generalize was measured. The generalization score reflects the ability of a trained network to generate the same behavior in response to a pair of odors (one odor was used during training, the other was not): a score of 1 means that the network generated the same behavior to both odors and a score of 0 means that the network generated opposite behaviors. (B) The generalization score averaged across pairs of similar odors (pairs with a cosine distance < 0.23) and across networks as a function of the number of odors used to trained a network is shown. (*n*=20) (C) The generalization score obtained for a pair of dissimilar odors (pairs with a cosine distance > 0.92) as a function of the number of odors used to train a network is shown. (*n*=20) Light grey line: B_0_G_0_ variant; blue line: B_1_G_0_ variant; green line: B_0_G_1_ variant; dark grey line: B_1_G_1_ variant; yellow line: C variant.

It is conceivable that group-structured networks are better at generalization because they generate a large number of overlapping representations. To test this idea, we trained different network variants with a number of odors, from 10 to 500. The odors used during the training step included odors that were found to generate pairs of representations with low cosine distances (bottom 2% of pairs for each network shown in Figure 8B); for each pair, only one of the two odors was used during the training step. After training, we tested the networks with the second odor of the pair. We considered that a network could generalize if the behavior triggered by the second odor was the same as the behavior it had generated with the first one. A value of 1 indicates that a network successfully generalized all the odors tested, whereas a value of 0.5 indicates that a network could not generalize and performed just as poorly as if it had randomly selected between the two possible behaviors. When we compared the ability of the different network variants to generalize odor pairs of low cosine distance, we obtained the highest generalization scores with the B_1_G_1_ variant and the lowest generalization scores with the B_0_G_0_ variant, regardless of the number of odors used during the training step (Figure 8B). The ability of a network to generalize is mostly driven by group structure, as the B_0_G_1_ variant shows higher generalization scores than the B_1_G_0_ variant. The C variant performs in a similar fashion as the B_1_G_0_ variant: both variants can generalize better than the B_0_G_0_ variant but not as well as the B_0_G_1_ and B_1_G_1_ variants. When we performed the same analysis using odor pairs with high cosine distances (top 2% of pairs for each network shown in Figure 8B), we failed to detect a similar phenomenon: instead, all variants performed the same and showed a generalization score of about 0.5 (Figure 8C).

There is a well-known tradeoff between generalizability and discriminability, notably in networks that encode stimuli as sparse activity patterns such as the ones analyzed here. Although networks that encode information sparsely are far better at discriminating, they are weaker at generalizing (Spanne and Jörntell, 2015). Thus, group-structured networks resolve this trade-off by gaining the ability to generalize across highly similar odors — here defined as odors activating PNs from the same group — while retaining the ability to discriminate by also producing highly dissimilar odors. Also, it is worth noting is that many network features — the ability of the group-structured networks to generate a wider range of representation similarities and the high median cosine distance produced by the bias-structured networks — were not detected by simply looking at their dimensionality. Thus, dimensionality alone does not capture all aspects of separability.

Altogether, these results suggest that networks built using groups can generate a wider range of odor representation similarities, including a large number of either highly overlapping or largely non-overlapping representations; this feature enables group-structured networks to generalize across certain odors while maintaining, overall, high discriminability.

## DISCUSSION

In this study, we devised a minimal model of the *Drosophila melanogaster* mushroom body — a cerebellum-like structure — that simulates odor representation and learning. We investigated how large-scale architectural features can instruct and constrain the ability of the mushroom body to process sensory information. We generated ten variants of this network that each differ in their degree of biased and grouped connectivity. One of these variants — the C variant — replicates the mushroom body input connections that were identified in the most recent *Drosophila melanogaster* connectome (Li et al., 2020a). The odor representations generated by the ten different network variants were analyzed quantitatively and qualitatively, and the ability of the network variants to form and recall associations was compared. First, we show that biases and groups reduce the overall dimensionality of the representation space generated by a network and therefore limit the number of associations it can recall accurately in a learning task. Second, we demonstrate that biases prioritize the robust representation of a subset of mono-odors — facilitating the formation and recall of associations made with these odors — compared to the representation of another subset of mono-odors, for which the formation and recall of associations is essentially disabled. Finally, we show that groups enhance the ability of a network to generate a wider range of representation similarities, enabling a network to generalize across similar odors while maintaining its ability to discriminate dissimilar odors.

### Assumptions and predictions

Our network was built using parameter values that have been derived experimentally whenever available, and when needed, parameter values that were fitted to the performance of the network. The latter set of parameters — for instance, the activation threshold of KCs and the *α*_LTD_ and *α*_LTP_ values used during the recall task — were fitted such that the network would generate an output that was similar to experimentally measured values. For instance, the relative strength of the feedforward excitation mediated by PNs and the global feedback inhibition mediated by APL was adjusted such that on average between 5 and 15% of the KCs would be activated by multi-odors, as has been measured experimentally (Honegger et al., 2011; Lin et al., 2014). Likewise, the *α*_LTD_ and *α*_LTP_ values were set such that *α*_LTD_ was greater than *α*_LTP_ as has been demonstrated experimentally (Cohn et al., 2015). Naturally, experimentally derived parameters, although based on available data, might not fully recapitulate the biology of the *Drosophila melanogaster* mushroom body and are contingent on the accuracy of the approaches used to measure them. For instance, the experimental data used to design the two different connectivity patterns — biases and groups — were derived from different experimental studies. We designed biases based on the non-uniform distribution of Kenyon cells inputs obtained in a study that mapped a subset of connections between projection neurons and Kenyon cells (Caron et al., 2013). We designed groups based on two different studies that reconstructed projection neurons and predicted their connectivity to Kenyon cells based on anatomical overlap (Jefferis et al., 2007; Lin et al., 2007).

Recently, the full connectome of the *Drosophila melanogaster* hemibrain was revealed, including all connections in the mushroom body (Li et al., 2020a). In this newly available data set, the connections between the projection neurons and Kenyon cells follow a non-uniform distribution of input similar to that measured in Caron et al., 2013, corresponding to our B_1_G_0_ network variant. Indeed, in almost all the tests we carried out, the C variant performed most similarly to the B_1_G_0_ variant. Likewise, group structure is less prevalent than originally thought (Jefferis et al., 2007; Lin et al., 2007). Careful analyses of the connectome data revealed a more subtle group pattern: a subset of 10 different types of projection neuron were found to connect more frequently to the same Kenyon cells, but these Kenyon cells do not integrate input exclusively from this group (Zheng et al., 2020). This structural feature of the *Drosophila melanogaster* mushroom body is most similar to our G_0.5_ variant. Interestingly, strong group structure has been found for other, non-olfactory projection neurons: two populations of Kenyon cells exclusively integrate input from projection neurons processing visual information (Li et al., 2020a, 2020b; Vogt et al., 2014, 2016). Although the set of experimentally-derived and fitted parameters may not perfectly replicate all the features of the mushroom body — and as such might limit the predictive scope of our study — we believe that, qualitatively, the functions enabled by biases and groups do realistically reflect actual biological functions. Below, we detail and discuss predictions emerging from our results.

### Prediction: biases prioritize the representation of a few ethologically pertinent stimuli

Several anatomical studies — including the recent hemibrain connectome — have revealed that the connections between projection neurons and Kenyon cells are biased. Interestingly, the projection neurons whose connections are most biased relay information from olfactory sensory neurons that are narrowly tuned to a few, specific odors that either elicit innate behaviors or that are used in various learned behaviors. Among the projection neurons forming the largest number of connections with Kenyon cells are neurons tuned to pheromones and food odors, including the DC3 projection neurons modeled as PN_10_ in this study. Conversely, among the projection neurons forming the smallest number of connections with Kenyon cells are neurons tuned to odors that elicit strong innate repulsion, including the VL1 projection neurons modeled as PN_40_ in this study. Such innate olfactory-driven behaviors are thought to be driven by the lateral horn, the other brain area innervated by olfactory projection neurons, and might not require the mushroom body. This idea remains to be tested experimentally. Therefore, biases might serve to shape the representation space of the mushroom body such that pertinent chemosensory information is prioritized: odors that need to be learned in a variety of contexts might be assigned a relatively larger portion of the space, whereas odors that ought not to be learned would occupy a smaller portion of that space. We predict that such prioritization of certain classes of stimuli, derived from our study of the numerically simple mushroom body, might also exist in the numerically more complex networks of vertebrates. For example, the input frequency of the granule cells of the vertebrate cerebellum — the equivalent of the Kenyon cells — might receive biased input, and these biases might enhance or suppress different sensory channels to reflect learning priorities.

### Prediction: groups enable networks to generalize stimuli with a similar meaning

In-depth analyses of two different connectomes have confirmed that the connections between projection neurons and Kenyon cells are not only biased, but also show some level of group structure: some projection neurons tend to connect more frequently to the same Kenyon cells than expected if these connections were completely random (Zheng et al., 2020). Although subtle most Kenyon cells integrate input broadly — this connectivity pattern is significant. Interestingly, the projection neurons identified to be part of this group — namely the VM3, DL2d, VA2, VA4, DP1m, DM1, DM2, DM3, DM4 and VM2 projection neurons — all receive input from olfactory sensory neurons broadly tuned to chemicals produced by fermenting fruits, the preferred food of *Drosophila melanogaster*. Similar to our results, the authors found that this low-level group structure enable the mushroom body to perform better in a classification task. Food odors — because they activate projection neurons that belong to the same group — generate largely overlapping representations in the mushroom body that can be used to form generalizable associations. The ability to generalize across odors with a similar meaning is arguably an advantageous feature for an associative brain center such as the mushroom body. For instance, upon associating the odor of one particular fermenting fruit with a nutritive reward, a fly will be able to learn that similar, albeit chemically different, odors released by fermentation of different fruits may also be worthwhile food sources. This strategy would enable a fly to exploit a wider range of resources.

Although advantageous for generalization, group structure is costly: our analyses clearly show that the dimensionality of the representation space of group-structured networks is drastically reduced when compared to the dimensionality of the space generated by biased or unstructured networks. This finding is in line with previous studies showing that there is a clear tradeoff between the ability of a network to generalize and its ability to discriminate (Spanne and Jörntell, 2015). Because the mushroom body shows overall a low level of group structure, it circumvents this trade-off in an efficient manner: the Kenyon cells that are more likely to receive convergent input from the projection neurons activated by food odors might be involved in the formation of highly generalizable associations, while the remaining Kenyon cells might be involved in the formation of more discriminable associations. Interestingly, such group structure exists in the vertebrate cerebellum, which clearly displays a general somatotopic organization (D’Angelo, 2018). Based on our findings, we predict that the generalization function enabled by group-structure is important for the learning function of the cerebellum, a brain center involved in movement coordination: similar types of movement or movements relating to a specific body part might be easier to generalize whereas different types of movements — reaching or walking for instance — might be easier to discriminate.

### Reaching optimality in cerebellum-like networks

Cerebellum-like structures are widespread: they can be found in the brains of vertebrates and invertebrates, even occurring in different centers of the same brain (Bell et al., 2008; Montgomery et al., 2012). Their occurrence in diverse brains suggests that the network architecture of cerebellum-like structures is well suited for associative learning. Conceptually, the high dimensionality enabled by completely random input connectivity would benefit the learning function of any cerebellum-like structure. We chose to focus our study on the extensively studied mushroom body in the context of the chemosensory ecology particular to *Drosophila melanogaster* because of the wealth of experimental data available. Our study clearly suggests that the mushroom body is not built using a one-fits-all design principle but rather built to reflect the expected ecology and learning priorities of a species. Although Kenyon cell input is largely unstructured, we found that specific connectivity patterns shape the representation space of the mushroom body such that olfactory information can be processed and used efficiently. While the specific connectivity patterns explored in our study are well-suited for the lifestyle of *Drosophila melanogaster*, they might not be as well-suited for other, closely related species with a different sensory ecology. Likewise, not all cerebellum-like structures process olfactory information — the vertebrate cerebellum processes primarily sensorimotor information — and, because connectomes are not yet available for these structures, it remains unclear whether biases and groups are used to shape representation space in the cerebellum in a similar manner as they do in the mushroom body. Altogether, we believe this study will pave the way toward understanding how ecological and evolutionary pressures shape cerebellum-like structures, enabling these structures to reach optimality by fine-tuning the way they represent and learn meaningful sensory information.

## AKNOWLEDGMENTS

We thank Ashok Litwin-Kumar, Florian Maderspacher and members of the Caron and Borisyuk laboratories for comments on the manuscript. This work has been funded in part by grants from the National Institute for Neurological Disorders and Stroke (R01 NS 106018, R01 NS 107970, R01 NS 109979) and the National Science Foundation (DMS 1853673 and IOS 2042397). Further financial support was provided by the Georges S. and Dolores Eccles Foundation (SJCC). Finally, the support and resources from the Center for High Performance Computing at the University of Utah are gratefully acknowledged.

## MATERIALS AND METHODS

### Mathematical equations used to build all network variants

#### First and second layer of the network: OSNs and PNs

We devised a four-layer network that simulates odor representation in the mushroom body. The first two layers of the network were based largely on previous models of the *Drosophila melanogaster* antennal lobe (Luo et al., 2010; Olsen et al., 2010). The first layer consists of 50 OSNs and connects in a one-to-one manner to the second layer of the network, which consists of 50 PNs. Each PN receives feedforward excitation from one OSN whose activity was based on available experimental data (see Realistic and expanded odor panels). Each PN is subject to lateral inhibition from a local inhibitory neuron (iLN) replicating available experimental data (Hong and Wilson 2013). The activity of PN *i* to odor *j* was generated using the equation:

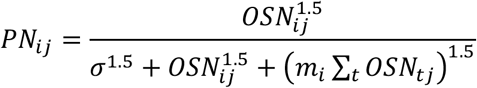

where *OSN_ij_* is the response of OSN *i* to odor *j* (firing frequency in Hz), *σ* = 12 is a constant fit to data. The inhibition is modeled as implicit, as represented by the last term in the denominator. Note that each PN has a different sensitivity to lateral inhibition, denoted by *m_i_*. Half of the 50 *m_i_*, values were experimentally determined while the remaining values were drawn from a normal distribution with the same mean and standard deviation as the experimentally-measured set (Hong and Wilson, 2015). This was repeated for each network realization.

#### Third layer of the network: KCs

The third layer consists of 2000 KCs. The activity of KC *k* to odor *j* was generated using the equation:

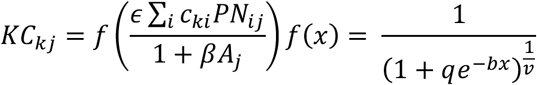

where *c_ki_* is the number of connections between PN *i* and KC *k*, *A_j_* is the response of the APL neuron to odor *j*, 2 and 4 control the strength of excitation from PNs and inhibition from APL, respectively (2 = 0.8, 4 = 0.03), and *f(x)* is a sigmoidal KC input-output function. All the PN–KC connections have the same strength. The parameters of *f(x)* were tuned such that approximately two inputs from active PNs were required to activate a KC and the resulting function was very steep but continuous (*q=20*, *b=32*, *v=10^-13^*). The activity (mV) of APL was generated using the equation:

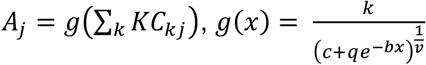

where the parameters of *g(x)* were fit to peak EPSP amplitudes evoked by extracellular stimulation of KCs (c=1.0653, q = 0.3847, b = 0.0379, v = 0.0884, k = 37.5910) (Papadopoulou et al., 2011). Thus, the resulting model of KCs and APL responses to an odor is a system of 2001 algebraic equations, which were solved using MATLAB’s *fsolve* command.

#### Fourth layer of the model: MBONs

The fourth layer consists of 2 MBONs, MBON+ and MBON-. See the “Recall capacity” section for a description of the equations used to simulate responses in MBON+ and MBON-.

### Connectivity patterns structuring PN–KC connections

We built nine different network variants by structuring the connections between PNs and KCs. In all network variants, the number of PNs connecting to a given KC is an integer drawn from a normal distribution — with a mean of seven and a standard deviation of 1.77 — that replicates features of Kenyon cells as measured experimentally (Caron et al., 2013). In the B_0_G_0_ network variants, each KC receives connections from a randomly selected group of PNs; in this case, the rates at which individual PNs connect with KCs follow a uniform distribution. In the B_1_ network variants, the uniform distribution of KC input was replaced with an experimentally derived non-uniform distribution (Caron et al. 2013). In the B_0.5_ network variants, the deviation of the non-uniform distribution from the uniform one was reduced by half. In the G_1_ network variants, PNs were divided into five groups based on two experimental data sets (Jefferis et al., 2007; Lin et al., 2007). Group 1 consists of PN_19_, PN_22_, PN_25_, PN_35_, PN_44_ and PN_45_ which correspond to the DM2, DM5, DP1m, VA7m, VM2 and VM3 projection neurons, respectively; group 2 consists of PN_27_, PN_34_, PN_37_ and PN_42_ which correspond to the VA1d, VA7l, VC2, and VL2p projection neurons, respectively; group 3 consists of PN_3_, PN_15_ and PN_50_ which correspond to the DA1, DL3 and VM7 projection neurons, respectively; group 4 consists of PN_1_, PN_2_, PN_9_, PN_10_, PN_12_, PN_23_ and PN_28_ which correspond to the 1, D, DC2, DC3, DL1, DM6 and VA1v projection neurons, respectively. There were no available data for the remaining 30 projection neurons and therefore the corresponding PNs were randomly assigned to a group such that each group comprises a total of 10 PNs. Groups 5 consists of 10 of these PNs. In the G_0.5_ network variants, only half of the KCs receive group-structure input whereas the other half receive unstructured input.

### Realistic and expanded odor panels

We built two different odor panels: a realistic odor panel — which contain 116 odors (110 multi- odors and 6 mono-odors) — and an expanded odor panel — which contain 2000 odors. In these panels, each odor elicits responses across all 50 OSNs. The realistic odor panel was built using an experimental data set in which the responses elicited in 24 different types of olfactory sensory neuron to 110 odors were recorded (Hallem and Carlson, 2006). The 110 multi-odors were simulated based on this data set: for each multi-odor, we assigned a firing rate to the 24 corresponding OSNs reflecting the experimental values; we simulated the missing responses — the responses elicited by each of the 110 odors in the remaining 26 olfactory sensory neurons — by resampling, randomly and with replacement, from the known responses recorded for the same odor. We executed this resampling step for each network realization. We added to these 110 multi-odors six mono-odors that elicit responses in only one type of OSN. Mono-odors were created based on different experimental studies that showed that some molecules — namely *cis-* vaccenyl acetate, geosmin, valencene, farnesol, CO_2_, and pyrrolidine — activate one type of olfactory sensory neuron exclusively or predominantly (Dweck et al., 2013; Kurtovic et al., 2007; Ronderos et al., 2014; Schlief and Wilson, 2007; Stensmyr et al., 2012; Suh et al., 2004). For each mono-odor, we simulated the responses of the corresponding OSN — namely OSN_3_, OSN_4_, OSN_8_, OSN_10_, OSN_26_ and OSN_40_ — based on the experimental values while maintaining the responses of that OSN to zero for all other odors, reflecting the specificity of these molecules for their cognate receptors. Thus, the realistic odor panel is a matrix with 50 OSNs responses to a total of 116 odors. The expanded odor panel was also built using the Hallem and Carlson data set (Hallem and Carlson, 2006). To this end, we generated 110 multi-odors as described above, and added 1890 simulated multi-odors. The OSN responses for the 1890 additional multi-odors were simulated by resampling, randomly and with replacement, from the responses reported in the Hallem and Carlson data set. We set the responses of OSN that respond to mono-odors — namely OSN_3_, OSN_4_, OSN_8_, OSN_10_, OSN_26_ and OSN_40_ — to zero for the reasons stated above. Thus, the realistic odor panel is a 50 by 116 matrix whereas the expanded odor panel is a 50 by 2000 matrix.

### Noise in the OSN layer

Gaussian noise was added to the OSN layer using two different equations. First, we added noise to the OSNs that were activated by the odor *j* using the equation:

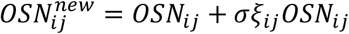

where *OSN_ij_* is the response of OSN *i* to odor *j*, *σ* ∈ [0,1] describes the overall level of noise and *ξ_ij_* is a random number drawn from a standard normal distribution. Second, we added noise to the OSNs that were not activated by odor *j* using the equation:

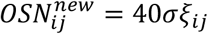

where 40 is a baseline amount of additional noise and *σ*, *ξ_ij_* are as described above. All responses that became negative were set to zero.

### Dimensionality

We estimated the dimension of the KC responses to an odor panel X — a matrix with 2,000 columns, each corresponding to a KC and a number of rows, each corresponding to an odor — by first computing the covariance of the given matrix C and, similar to previous studies (Litwin-Kumar et al. 2017), we estimated the dimension using the equation:

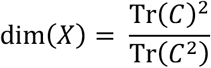

where Tr(*C*) = ∑_*i*_ λ_*i*_ and {λ_*i*_} is the set of eigenvalues of C. Finally, dimensions were normalized by the highest dimension found across network variants and trials.

### Recall capacity

#### Network training and testing

We tested the ability of a network to form and recall associations based largely on a previous study (Peng and Chittka 2017). Each MBON initially receives feedforward excitation from all KCs with equal strength. One MBON, MBON+, encodes attraction while the other MBON, MBON-, encodes aversion. The activity of each MBON was generated using the following equations:

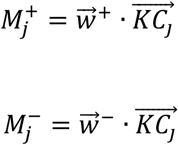

where 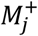 and 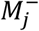 are the responses of MBON+ and MBON- to odor *j*, respectively, 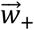 and 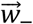 are vectors describing the weight of connections from each Kenyon cell to MBON+ and MBON-, and 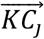 is a vector describing the responses of all Kenyon cells to odor *j*.

During the training step, we subjected the KC to MBON connections to an anti-Hebbian learning rule: the connections between the KCs activated by an odor and the MBON encoding the opposite behavioral outcome were weakened by an *α*_LTD_ factor while the connections between the inactive KCs and the same MBON were strengthened by an *α*_LTP_ factor (Cohn et al., 2015; Peng and Chittka 2017). This process can be summarized by the following set of equations.

Initially, the weight of each connection is set to 1 and if a positive behavioral outcome is paired with the *j*^th^ KC odor representation, then:

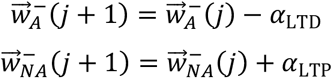

where *A* and *NA* are the indices of active and inactive KCs respectively and *α*_LTP_ = 0.25 and *α*_LTP_ = 0.01 are parameters describing the strength of *α*_LTD_ and *α*_LTP_, respectively. Similarly, if a negative behavioral outcome is paired with the *j*^th^ KC odor representation, then:

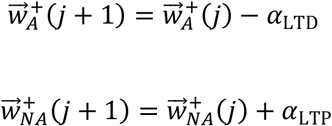

Throughout the training step, the connections weights were constrained to lie on the interval [0,1.5], and were capped at the ends of this interval. Each network was trained with *n* odors (*n*=20, 40, 60, …, 1200) and the behavioral outcome were randomly assigned to an odor.

Once the feedforward networks 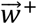 and 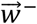 were trained, the training step ended, and the testing step started. During the testing step, the connections were static, the odors used during the training step were presented to the trained network sequentially, and the model classified the KC odor representations as attractive or aversive by assigning it the valence of the most active MBON. We defined the recall capacity as the number of odors that a trained network could classify correctly with 75% accuracy. Finally, we examined the relative importance of *α*_LTD_ and *α*_LTP_ during the testing step by holding *α*_LTD_ constant at 0.25 and varying *α*_LTP_ cross 50 values spaced evenly in a logarithmic scale between 0.25 and 0.25 ∗ 10^-7^and repeating the recall capacity procedure described above for each pair of *α*_LTD_, *α*_LTP_ values.

#### UMAP dimensionality reduction algorithm

We performed dimension reduction on the KC representations — from 2000 to 2 dimensions — for visualization and clustering. To this end, we used the nonlinear dimension reduction software UMAP (McInnes et al., 2018). In short, UMAP finds a low dimensional manifold on to which high dimensional data can be projected using topological data analysis techniques. There are four main hyperparameters that determine the output of UMAP: the number of nearest neighbors used, the minimum Euclidean distance between data points that is allowed in the low dimensional representation, the dimension of the low dimensional representation and the distance measure that is used. Throughout our study we used 25 nearest neighbors, a minimum distance of 0.1, an output dimension of 2 and cosine distance to measure distances.

#### K-means clustering tool, clustering accuracy and clustering errors

The KC representations generated for 12 odors (two mono-odors and ten multi-odors) at various noise levels were reduced using the UMAP algorithm; the resulting representations were analyzed using the k-means clustering tool available in MATLAB. We searched for 12 clusters, one cluster for each odor. Because the cluster centroids were seeded randomly, we applied the *k-means* clustering tool to each set of UMAP representations ten times.

We used the adjusted Rand index to measure the clustering accuracy of the k-means tool. If a set of *n* data points is divided in to the following clusters {*X*_1_, *X*_2_, …, *X*_*k*_} and also has ground truth labels {*Y*_1_, *Y*_2_, …, *Y*_*k*_} then one can construct a contingency table whose entries *n_ij_* are the sizes of the intersections of *X_i_* and *Y_i_*, *a_i_* is the sum of row *i*, and *b_i_* is the sum of column *j*. The adjusted Rand index is a measure of the similarity of the clustering to the ground truth labels that accounts for random intersections (Hubert and Arabie, 1985). We measured the adjusted Rand index using the following equations:

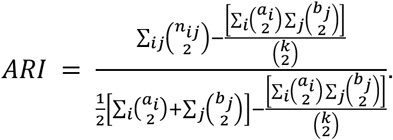

We calculated the clustering errors for the representations generated for each odor by finding the largest number of representations that were clustered together (out of the possible ten representations) and subtracting that number from ten. The result is the number of representations that were not clustered with the “main” cluster. For example: if one cluster consisted of seven representations of the same odor, the clustering error would be three.

#### Cosine distance

We calculated the cosine distance between pairs of Kenyon cell odorant representations using the following equation:

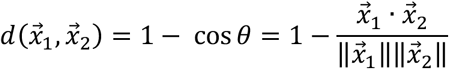

where 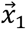 and 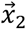 are vectors of Kenyon cell odorant representations with the entries of each vector corresponding to the activity level of the Kenyon cell with the same index and f is the angle between the two vectors.

## COMPETING INTERESTS

The authors declare that no competing interests exist.

---

**Supplementary Figure 1.**
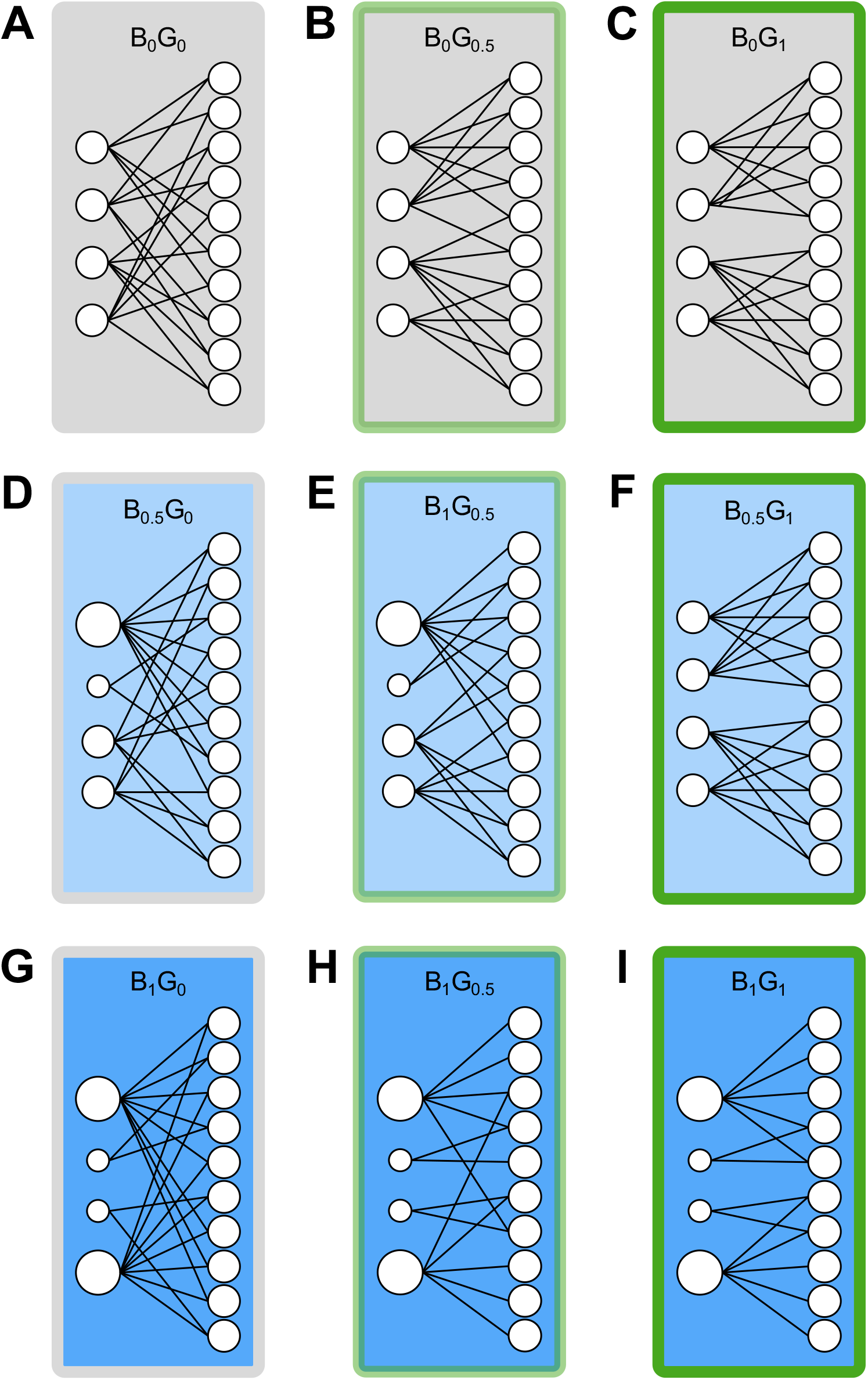
The network variants were built using different degrees of structure. Nine network variants were generated, each built using biases, groups, a combination of both connectivity patterns, or no structure at all. The B_0_G_0_ network variant is completely unstructured (A; grey fill and grey outline). The G_0.5-1_ network variants were built using groups: either half of the KCs (B,E,H; light green outline) or all the KCs (C,F,I; dark green outline) are group-structured. The B_0.5-1_ network variants were built using biases: the number of connections formed by a given PN was assigned using either a uniform distribution of input (A-C; light grey fill), the non-uniform distribution of input measured experimentally (G-I; dark blue fill) or a value halfway in between (D-F; light blue fill).

**Supplementary Figure 2.**
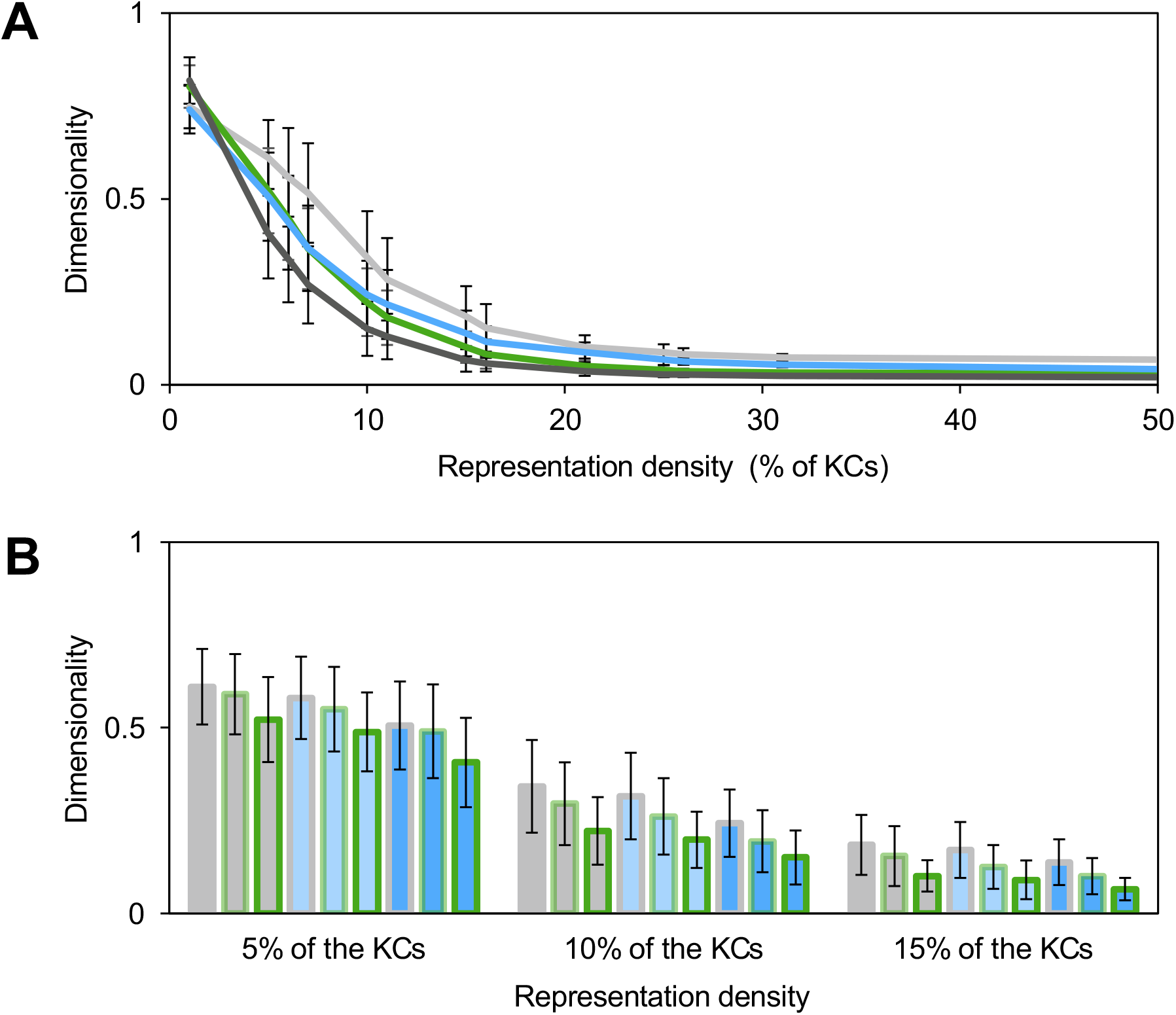
Structure reduces dimensionality when the representation density is fixed. (A) Each of the ten network variants were used to generate 5,000 KC representations using the expanded odor panel; the representation density was fixed at values ranging from 1 to 50% of the KCs; the relative dimensionality of the resulting 5,000 representations was measured and averaged across network realizations (*n*=20). Error bars represent the standard deviation. Light grey line: B_0_G_0_ networks; blue line: B_1_G_0_ networks; green line: B_0_G_1_ networks; dark grey line: B_1_G_1_ networks. (B) The average dimensionality obtained for realistic representations densities (at 5, 10 and 10% of the KCs) across all nine network variants are shown. Error bars represent the standard deviation. Grey fill and outline: B_0_G_0_ variant; blue fill and grey outline: B_1_G_0_ variant; grey fill and green outline: B_0_G_1_ variant; blue fill and green outline: B_1_G_1_ variant.

**Supplementary Figure 3.**
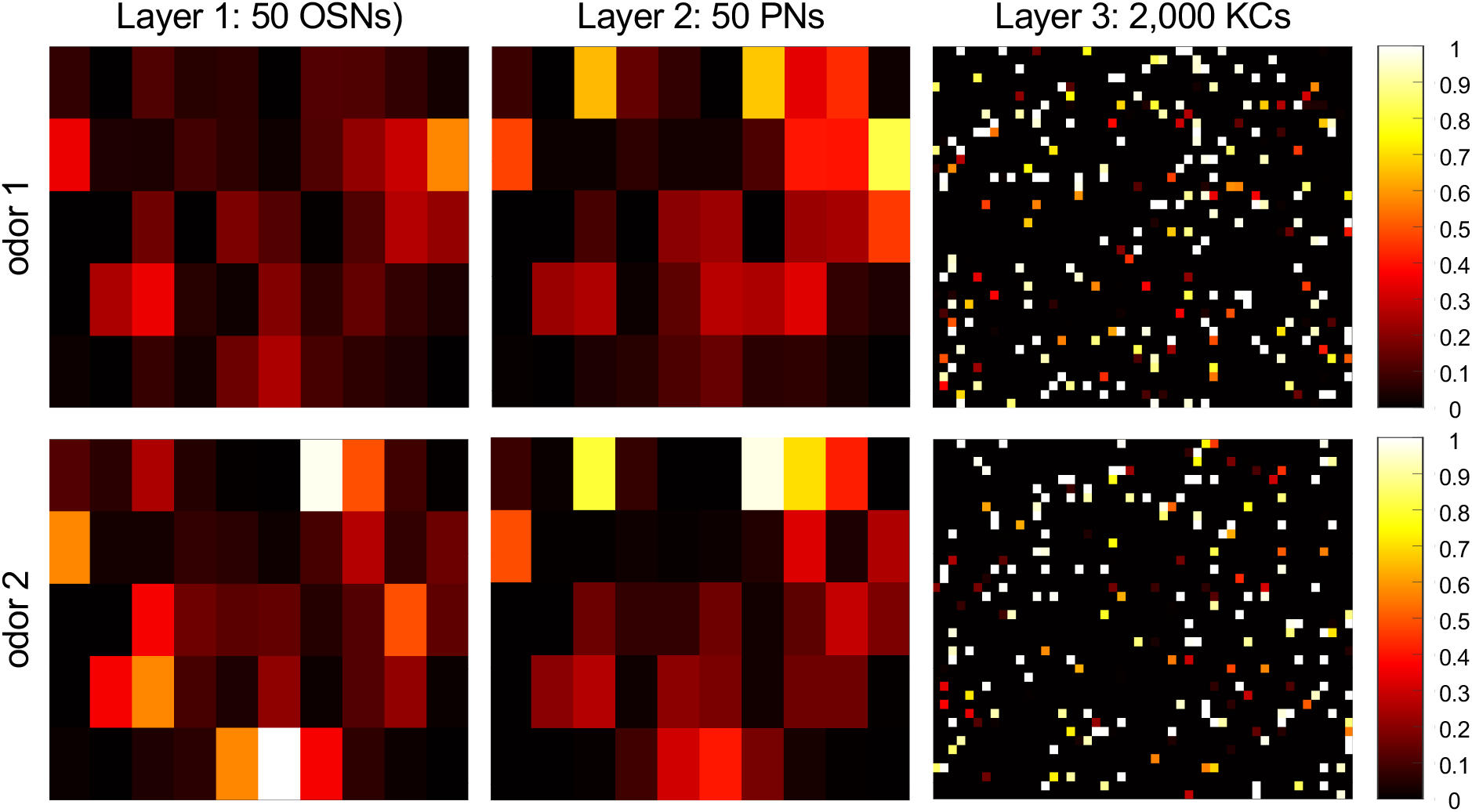
Odor representations in the first three layers of the network. Odors from the realistic odor panel (odor 1: -(trans)-caryophyllene; odor 2: p-cymene) activate a number of OSNs at different rates; OSNs activity patterns are first transformed into PN activity patterns and then in KC activity patterns.

## Notes

### Competing Interest Statement

The authors have declared no competing interest.

## REFERENCES

Albus, J. (1971). A theory of cerebellar function. Math. Biosci. 10, 25–61.

Aso, Y., Sitaraman, D., Ichinose, T., Kaun, K.R., Vogt, K., Belliart-guérin, G., Plaçais, P., Robie, A.A., Yamagata, N., Schnaitmann, C., et al. (2014a). Mushroom body output neurons encode valence and guide memory-based action selection in Drosophila. 1–42.

Aso, Y., Hattori, D., Yu, Y., Johnston, R.M., Iyer, N.A., Ngo, T.T.B., Dionne, H., Abbott, L.F., Axel, R., Tanimoto, H., et al. (2014b). The neuronal architecture of the mushroom body provides a logic for associative learning. Elife 3, e04577.

Babadi, B., and Sompolinsky, H. (2014). Sparseness and Expansion in Sensory Representations. Neuron 83, 1213–1226.

Bell, C.C., Han, V., and Sawtell, N.B. (2008). Cerebellum-like structures and their implications for cerebellar function. Annu. Rev. Neurosci. 31, 1–24.

Bhandawat, V., Olsen, S.R., Gouwens, N.W., Schlief, M.L., and Wilson, R.I. (2007). Sensory processing in the Drosophila antennal lobe increases reliability and separability of ensemble odor representations. Nat. Neurosci. 10, 1474–1482.

Caron, S.J.C., Ruta, V., Abbott, L.F., and Axel, R. (2013). Random convergence of olfactory inputs in the Drosophila mushroom body. Nature 497, 113–117.

Cayco-Gajic, N.A., and Silver, R.A. (2019). Re-evaluating Circuit Mechanisms Underlying Pattern Separation. Neuron 101, 584–602.

Cayco-Gajic, N.A., Clopath, C., and Silver, R.A. (2017). Sparse synaptic connectivity is required for decorrelation and pattern separation in feedforward networks. Nat. Commun. 8, 1–11.

Cohn, R., Morantte, I., Cohn, R., Morantte, I., and Ruta, V. (2015). Coordinated and Compartmentalized Neuromodulation Shapes Sensory Processing in Sensory Processing in Drosophila. Cell 163, 1742–1755.

D’Angelo, E. (2018). Physiology of the cerebellum. In Handbook of Clinical Neurology, (Elsevier B.V.), pp. 85–108.

Dweck, H.K.M., Ebrahim, S.A.M., Kromann, S., Bown, D., Hillbur, Y., Sachse, S., Hansson, B.S., and Stensmyr, M.C. (2013). Olfactory preference for egg laying on citrus substrates in Drosophila. Curr. Biol. 23, 2472–2480.

Eichler, K., Li, F., Litwin-Kumar, A., Park, Y., Andrade, I., Schneider-Mizell, C.M., Saumweber, T., Huser, A., Eschbach, C., Gerber, B., et al. (2017). The complete connectome of a learning and memory centre in an insect brain. Nature 548, 175–182.

Farris, S.M. (2011). Are mushroom bodies cerebellum-like structures? Arthropod Struct. Dev. 40, 368–379.

Fusi, S., Miller, E.K., and Rigotti, M. (2016). Why neurons mix: High dimensionality for higher cognition. Curr. Opin. Neurobiol. 37, 66–74.

Gruntman, E., and Turner, G.C. (2013). Integration of the olfactory code across dendritic claws of single mushroom body neurons. Nat. Neurosci. 16, 1821–1829.

Hallem, E.A., and Carlson, J.R. (2006). Coding of Odors by a Receptor Repertoire. Cell 125, 143–160.

Hige, T., Aso, Y., Rubin, G.M., and Turner, G.C. (2015). Plasticity-driven individualization of olfactory coding in mushroom body output neurons. Nature 526, 258–262.

Honegger, K.S., Campbell, R.A.A., and Turner, G.C. (2011). Cellular-resolution population imaging reveals robust sparse coding in the drosophila mushroom body. J. Neurosci. 31, 11772–11785.

Hong, E.J., and Wilson, R.I. (2013). Olfactory neuroscience: Normalization is the norm. Curr. Biol. 23, R1091–R1093.

Hong, E.J., and Wilson, R.I. (2015). Simultaneous Encoding of Odors by Channels with Diverse Sensitivity to Inhibition. Neuron 85, 573–589.

Hubert, L., and Arabie, P. (1985). Comparing Partitions. J. Classif. 218, 193–218.

Huerta, R., Nowotny, T., García-Sanchez, M., Abarbanel, H.D.I., and Rabinovich, M.I. (2004). Learning classification in the olfactory system of insects. Neural Comput. 16, 1601–1640.

Jefferis, G.S.X.E., Potter, C.J., Chan, A.M., Marin, E.C., Rohlfing, T., Maurer, C.R., and Luo, L. (2007). Comprehensive Maps of Drosophila Higher Olfactory Centers: Spatially Segregated Fruit and Pheromone Representation. Cell 128, 1187–1203.

Jortner, R.A., Farivar, S.S., and Laurent, G. (2007). A simple connectivity scheme for sparse coding in an olfactory system. J. Neurosci. 27, 1659–1669.

Kurtovic, A., Widmer, A., and Dickson, B.J. (2007). A single class of olfactory neurons mediates behavioural responses to a Drosophila sex pheromone. Nature 446, 542–546.

Li, F., Lindsey, J., Marin, E.C., Otto, N., Dreher, M., Dempsey, G., Stark, I., Bates, A.S., Pleijzier, M.W., Schlegel, P., et al. (2020a). The connectome of the adult Drosophila mushroom body provides insights into function. Elife 9, e62576.

Li, J., Mahoney, B.D., Jacob, M.S., and Caron, S.J.C. (2020b). Visual Input into the Drosophila melanogaster Mushroom Body. Cell Rep. 32.

Lin, A.C., Bygrave, A.M., De Calignon, A., Lee, T., and Miesenböck, G. (2014). Sparse, decorrelated odor coding in the mushroom body enhances learned odor discrimination. Nat. Neurosci.

Lin, H.H., Lai, J.S.Y., Chin, A.L., Chen, Y.C., and Chiang, A.S. (2007). A Map of Olfactory Representation in the Drosophila Mushroom Body. Cell 128, 1205–1217.

Litwin-Kumar, A., Harris, K.D., Axel, R., Sompolinsky, H., and Abbott, L.F. (2017). Optimal Degrees of Synaptic Connectivity. Neuron 93, 1153–1164.e7.

Luo, S.X., Axel, R., and Abbott, L.F. (2010). Generating sparse and selective third-order responses in the olfactory system of the fly. Proc. Natl. Acad. Sci. 107, 10713–10718.

Marin, E.C., Jefferis, G.S.X.E., Komiyama, T., Zhu, H., and Luo, L. (2002). Representation of the glomerular olfactory map in the Drosophila brain. Cell 109, 243–255.

Marr, D. (1969). A Theory of Cerebellar Cortex. J. Physiol. 202, 437–470.

McInnes, L., Healy, J., and Melville, J. (2018). UMAP: Uniform Manifold Approximation and Projection for Dimension Reduction.

Montgomery, J.C., Bodznick, D., and Yopak, K.E. (2012). The cerebellum and cerebellum-like structures of cartilaginous fishes. Brain. Behav. Evol. 80, 152–165.

Münch, D., and Galizia, C.G. (2016). DoOR 2.0 - Comprehensive Mapping of Drosophila melanogaster Odorant Responses. Sci. Rep. 6, 1–14.

Murthy, M., Fiete, I., and Laurent, G. (2008). Testing Odor Response Stereotypy in the Drosophila Mushroom Body. Neuron 59, 1009–1023.

Olsen, S.R., and Wilson, R.I. (2008). Lateral presynaptic inhibition mediates gain control in an olfactory circuit. Nature 452, 956–960.

Olsen, S.R., Bhandawat, V., and Wilson, R.I. (2010). Divisive Normalization in Olfactory Population Codes. Neuron 66, 287–299.

Owald, D., and Waddell, S. (2015). Olfactory learning skews mushroom body output pathways to steer behavioral choice in Drosophila. Curr. Opin. Neurobiol. 35, 178–184.

Owald, D., Felsenberg, J., Talbot, C.B., Das, G., Perisse, E., Huetteroth, W., and Waddell, S. (2015). Activity of defined mushroom body output neurons underlies learned olfactory behavior in Drosophila. Neuron 86, 417–427.

Papadopoulou, M., Cassenaer, S., Nowotny, T., and Laurent, G. (2011). Normalization for sparse encoding of odors by a wide-field interneuron. Science (80-.). 332, 721–725.

Peng, F., and Chittka, L. (2017a). Report A Simple Computational Model of the Bee Mushroom Body Can Explain Seemingly Complex Forms of Olfactory Learning and Memory Report A Simple Computational Model of the Bee Mushroom Body Can Explain Seemingly Complex. Curr. Biol. 27, 224–230.

Peng, F., and Chittka, L. (2017b). A Simple Computational Model of the Bee Mushroom Body Can Explain Seemingly Complex Forms of Olfactory Learning and Memory. Curr. Biol. 27, 224– 230.

Ronderos, D.S., Lin, C.C., Potter, C.J., and Smith, D.P. (2014). Farnesol-detecting olfactory neurons in drosophila. J. Neurosci. 34, 3959–3968.

Sawtell, N.B., and Bell, C.C. (2008). Adaptive processing in electrosensory systems: Links to cerebellar plasticity and learning. J. Physiol. Paris 102, 223–232.

Scheffer, L., Xu, C.S., Januszewski, M., Lu, Z., Takemura, S., Hayworth, K., Huang, G., Shinomiya, K., Maitin-Shepard, J., Berg, S., et al. (2020). A connectome and analysis of the adult drosophila central brain. Elife 9, e57443.

Schlief, M.L., and Wilson, R.I. (2007). Olfactory processing and behavior downstream from highly selective receptor neurons. Nat. Neurosci. 10, 623–630.

Séjourn, J., Plaçais, P.Y., Aso, Y., Siwanowicz, I., Trannoy, S., Thoma, V., Tedjakumala, S.R., Rubin, G.M., Tchénio, P., Ito, K., et al. (2011). Mushroom body efferent neurons responsible for aversive olfactory memory retrieval in Drosophila. Nat. Neurosci. 14, 903–910.

Spanne, A., and Jörntell, H. (2015). Questioning the role of sparse coding in the brain. Trends Neurosci. 38, 417–427.

Stensmyr, M.C., Dweck, H.K.M., Farhan, A., Ibba, I., Strutz, A., Mukunda, L., Linz, J., Grabe, V., Steck, K., Lavista-Llanos, S., et al. (2012). A conserved dedicated olfactory circuit for detecting harmful microbes in drosophila. Cell 151, 1345–1357.

Stevens, C.F. (2015). What the fly’s nose tells the fly’s brain. Proc. Natl. Acad. Sci. 112, 9460– 9465.

Suh, G.S.B., Wong, A.M., Hergarden, A.C., Wang, J.W., Simon, A.F., Benzer, S., Axel, R., and Anderson, D.J. (2004). A single population of olfactory sensory neurons mediates an innate avoidance behaviour in Drosophila. Nature 431, 854–859.

Turner, G.C., Bazhenov, M., and Laurent, G. (2008). Olfactory representations by Drosophila mushroom body neurons. J. Neurophysiol. 99, 734–746.

Vogt, K., Schnaitmann, C., Dylla, K. V., Knapek, S., Aso, Y., Rubin, G.M., and Tanimoto, H. (2014). Shared mushroom body circuits underlie visual and olfactory memories in Drosophila. Elife 3, e02395.

Vogt, K., Aso, Y., Hige, T., Knapek, S., Ichinose, T., Friedrich, A.B., Turner, G.C., Rubin, G.M., and Tanimoto, H. (2016). Direct neural pathways convey distinct visual information to drosophila mushroom bodies. Elife 5, 1–13.

Wilson, R.I. (2013). Early Olfactory Processing in Drosophila: Mechanisms and Principles. Annu. Rev. Neurosci. 36, 217–241.

Wilson, R.I., Turner, G.C., and Laurent, G. (2004). Transformation of Olfactory Representations in the Drosophila Antennal Lobe. Science (80-.). 303, 366–370.

Wong, A.M., Wang, J.W., and Axel, R. (2002). Spatial representation of the glomerular map in the Drosophila protocerebrum. Cell 109, 229–241.

Zheng, Z., Lauritzen, J.S., Perlman, E., Robinson, C.G., Nichols, M., Milkie, D., Torrens, O., Price, J., Fisher, C.B., Sharifi, N., et al. (2018). A Complete Electron Microscopy Volume of the Brain of Adult Drosophila melanogaster. Cell 174, 730–743.e22.

Zheng, Z., Li, F., Fisher, C., Ali, I.J., Sharifi, N., Calle-Schuler, S.A., Hsu, J., Masoodpanah, N., Kmecova, L., Kazimiers, T., et al. (2020). Structured sampling of olfactory input by the fly mushroom body. BioRxiv.

